# Frequency-dependent interactions determine outcome of competition between two breast cancer cell lines

**DOI:** 10.1101/2020.03.06.979518

**Authors:** Audrey R. Freischel, Mehdi Damaghi, Jessica J. Cunningham, Arig Ibrahim-Hashim, Robert J. Gillies, Robert A. Gatenby, Joel S. Brown

## Abstract

Tumors are highly dynamic ecosystems in which diverse cancer cell subpopulations compete for space and resources. These complex, often non-linear interactions govern continuous spatial and temporal changes in the size and phenotypic properties of these subpopulations. Because intra-tumoral blood flow is often chaotic, competition for resources may be a critical selection factor in progression and prognosis. Here, we quantify resource competition using 3D spheroid cultures with MDA-MB-231 and MCF-7 breast cancer cells. We hypothesized that MCF-7 cells, which primarily rely on efficient aerobic glucose metabolism, would dominate the population under normal pH and low glucose conditions; and MDA-MB-231 cells, which exhibit high levels of glycolytic metabolism, would dominate under low pH and high glucose conditions. In spheroids with single populations, MCF-7 cells exhibited equal or superior intrinsic growth rates (density-independent measure of success) and carrying capacities (density-dependent measure of success) when compared to MDA-MB-231 cells under all pH and nutrient conditions. Despite these advantages, when grown together, MCF-7 cells do not always outcompete MDA-MB-231 cells. MDA-MB-231 cells outcompete MCF-7 cells in low glucose conditions and coexistence is achieved in low pH conditions. Under all conditions, MDA-MB-231 has a stronger competitive effect (frequency-dependent interaction) on MCF-7 cells than vice-versa. This, and the inability of growth rate or carrying capacity when grown individually to predict the outcome of competition, suggests a reliance on frequency-dependent interactions and the need for competition assays. We frame these results in a game-theoretic (frequency-dependent) model of cancer cell interactions and conclude that competition assays can demonstrate critical density-independent, density-dependent and frequency-dependent interactions that likely contribute to *in vivo* outcomes.

**Highlights:** Demonstrate how mixed-culture spheroids must be used to characterize competition between two cancer cell lines.

Competition alters growth dynamics of cancer cells.

Competition growth models can be used to quantify density-independent, density-dependent and frequency-dependent effects on competition.

Competition affects tumor progression and structure, making it key to understanding tumor development and evolution.

## INTRODUCTION

Each malignant tumor typically contains diverse and heterogeneous habitats due to spatial and temporal variations in cell density, immune infiltration, and blood flow (1–3). Here, we investigate the evolutionary effects of variations in substrate (e.g. glucose, oxygen, and glutamine) and metabolites (e.g. H^+^) that are typically governed by alterations in blood flow. These changes, in turn, can alter the local fitness (competitive advantage) of the extant cell populations that can have dramatic evolutionary effects on the phenotypic and genotypic properties of the cancer cells.

Recently, two general “niche construction” strategies were observed in clinical and pre-clinical cancers (4, 5). The first consists of cancer cells that possess high levels of aerobic glycolysis that generate an acidic environment, which promotes invasion and protects cells from the immune system. These cells are typically invasive and motile. In the second niche construction strategy, cancer cells promote growth by recruiting and promoting blood vessels. This angiogenic phenotype is typically non-invasive and maintains near-normal aerobic metabolism of glucose. These divergent phenotypes can be found within the same tumor (6–9). The invasive “pioneer” phenotype is optimally adapted to survive and thrive at the tumor edge. There it can co-opt normal tissue vasculature as it invades, and its acidic environment provides protection from immune cells at the tumor-host interface. In contrast, the “engineer” cancer cells are well suited to the interior regions of the tumor where their angiogenic phenotype can, with varying degrees of success, provide blood flow necessary to deliver substrate and remove metabolites. When the pioneers and engineers interact, they presumably compete for both space and substrate. However, the specific phenotypic traits that confer maximal fitness under different micro-environmental conditions have not been well established.

Competition between two species leads to four possible outcomes: 1) species A outcompetes B, 2) B outcompetes A, 3) A or B outcompetes the other depending upon who establishes first (alternate stable states), or 4) the two species coexist. These population dynamics can be mathematically framed through the Lotka-Volterra equations for inter-specific competition (10, 11). Derived independently by Lotka (1910) and Volterra (1920), the eponymous equations were first tested empirically by Gause (1932) in the, now classic, competition experiments using various species of Paramecium and yeast (12, 13). This, and subsequent work on fruit flies, flour beetles, and various natural systems, have confirmed the predictive power of the Lotka-Volterra equations (14, 15). Such assays utilize growth dynamics to elucidate the critical phenotypic properties that determine fitness under different environmental conditions (16).

In ecological systems, growth dynamics are divided into three categories: *density dependent* (DD), *density independent* (DI), and *frequency dependent* (FD) effects. Density dependent (DD) effects occur when one cancer cell type outcompetes another because it can achieve a higher equilibrium cell density (carrying capacity) under conditions of space or nutrient limitations. *Density independent* (DI) effects include intrinsic growth rate, which measures the capacity of a cell line to grow exponentially under ideal conditions. In cell biology, these (DD and DI) are often used as indicators of cell fitness (17–19). This, however, ignores the FD interactions, which often determine competitive outcomes. Frequency dependent (FD) effects occur when one cell type outcompetes another because its phenotype disproportionately depresses the growth rate of another as a result of more rapid resource uptake, or via direct interference with the other cell type. This game theoretic effect implies that the success of an individual cell is context dependent. It depends not only on the density of other cells around it, but the phenotypes of those cells as well (20–23). Because of this, FD effects and the outcome of competition can only be evaluated using Gause-style competition assays in which mixtures of the two cell types are grown together using different starting frequencies.

Here, we utilize Gause-style competition assays to investigate the competitive interaction between engineer and pioneer phenotypes. These strategies will be represented by the well-characterized MDA-MB-231 and MCF-7 cell lines, which are regularly used *in vitro* and *in vivo* as representative of highly glycolytic and invasive, and low glycolytic and non-invasive phenotypes, respectively (24–26). Any number of cancer cell lines would be good candidates for competition assays covering the myriad of potential factors that affect cellular growth and viability.

Assays were conducted in 3D spheroid cultures with varying starting frequencies of each cell type. The spheroid system was selected as an *in vitro* intermediate between 2D cell culture, which may reach an equilibrium state through confluence rather than resource competition, and *in vivo* subcutaneous mouse models, which provide less opportunity for replication and frequent monitoring of the two cell populations (27–34). Spheroids were grown in various combinations of pH and glucose concentrations, letting the 3D structure establish natural nutrient and oxygen gradients. Glutamine availability was also varied to test for growth and competition effects due to secondary nutrients.

As in the Gause experiments, measures of growth dynamics were used to evaluate competitive ability. DI, DD, and FD effects corresponded to selection for higher growth rates, carrying capacities, and competitive ability, respectively. Carrying capacities (DD) were estimated from monoculture spheroids under varying culture conditions. Initial growth rates (DI) of each cell type were estimated from both mono- and co-cultured spheroids. As FD effects rely on interactions between cell types, their values were estimated using the Lotka-Volterra competition model. In order to remain agnostic to the mode of tumor spheroid growth, our growth model was selected by first fitting monoculture spheroids to four common growth models (exponential, Monod, logistic, and Gompertz), then selecting the model which best approximated the growth of monoculture spheroids in all culture conditions. This model was then expanded to its two-species competition form and used to obtain estimates of competition coefficients for each cell line under eight different nutrient conditions.

We expected both cell types to have higher growth rates and carrying capacities at physiological pH (pH 7.4) and higher nutrient levels. MCF-7 cells, which predominantly use oxidative phosphorylation for metabolism of glucose, were predicted to be significantly inhibited by low pH, while MDA-MB-231 cells, which exhibit the Warburg phenotype with high levels of aerobic glycolysis, were expected to be more tolerant. Because the glycolytic MDA-MB-231 cells need more glucose to produce energy compared to the oxygen-consuming MCF-7 cells, we expected the former to respond more favorably to increased glucose concentrations than the latter. We also predicted that, based on their metabolic differences, MCF-7 cells would be fitter under conditions of normal pH and low glucose; and vise-versa for the MDA-MB-231 cells. Herein, we test our hypothesis through competition experiments between the MCF-7 and MDA-MB-231 cells under varying environmental conditions. We report the outcome of these competition assays as well as the effect of nutrient availability on DD and DI effects and evaluate their ability to indicate competitive outcome. We used fluorescent measurements to approximate cell growth in the spheroid and propose that the logistic growth equation best approximates the growth of both cell types in the majority of culture conditions. We then used Lotka-Volterra competition equations to determine the contribution of population growth rates (DI), maximal population size (DD) and/or competition coefficients (FD) on the outcome of interspecific competition. Lastly, we compared these results to the outcome of competition between MDA-MB-231 and MCF-7 from mixed tumors *in vivo*.

## RESULTS

### MDA-MB-231 generally out-compete MCF-7 cells under all pH, glucose and glutamine levels, and starting frequencies

Fluorescently labeled MCF-7 (GFP) and MDA-MB-231 (RFP) cells allowed us to non-destructively measure population dynamics within each micro-spheroid. We used this spheroid system to: 1) observe the outcome of competition (mixed culture), 2) compare population growth models (mono-cultures), 3) measure intrinsic growth rates (DI), 4) measure carrying capacities (DD), and 5) determine competition coefficients (FD). In experiments 1 and 2, spheroids started with a plating density of 20,000 and 10,000 cells, respectively, with initial conditions of 0%, 20%, 40%, 60%, 80% and 100% MDA-MB-231 cells relative to MCF-7 cells. In experiment 1, additional treatments included neutral (7.4) versus low (6.5) pH and varying levels of glucose (0, 1, 2, 4.5 g/L) (Figure 1B). Experiment 2 had treatments of pH (7.4, 6.5), glucose (0, 4.5 g/L), and glutamine positive and starved media (Figure 1C). The ranges of these values encompass both physiologic and extreme tumor conditions. For example, pH ~6.5 and very low levels of glucose have been measured near the necrotic core of the tumor. Conversely, high glucose and physiologic pH are present at the tumor edge and near vasculature, and also reflect standard culture conditions for our cell lines. Both experiments were further subdivided into parts A and B based on culture conditions (Figure 1). In experiment 1, microspheres were grown for 670 hours with fluorescence measured every 24-72 hours. In experiment 2, microspheres were grown for 550 hours with fluorescence measured every 6-12 hours. Growth medium was replaced every 4 days without disturbing the integrity of the microsphere.

**Figure 1:**
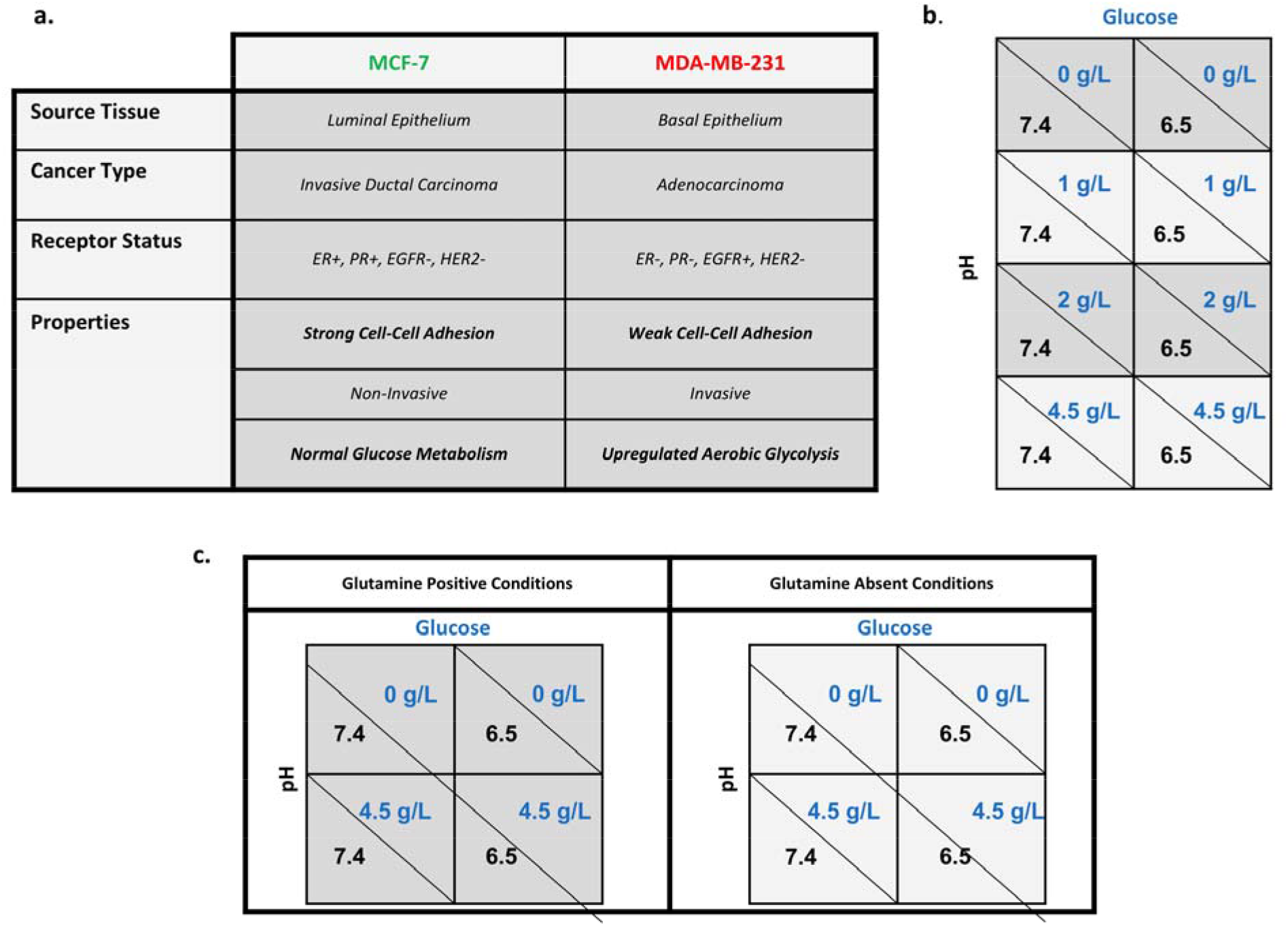
Cell Line Descriptions and Experimental Design A. Phenotype and source information for the “niche constructing” MCF-7 and the “pioneer” MDA-MB-231 cell lines. **B.** Growth dynamics of monoculture and mixed population tumor spheroids in glucose concentrations of 0 g/L and 2 g/L (in dark gray) and of 1 g/L and 4.5 g/L (in light gray) were compared at physiological (7.4) and low (6.5) pH. **C.** Experiment 2 measured the growth dynamics of mixed population tumor spheroids in 4.5 g/L glucose and 0 g/L glucose, in physiological (7.4) and low (6.5) pH, and with (in dark gray) and without (in light gray) glutamine to evaluate differences in growth due to the presence of a secondary nutrient.

Counter to our hypothesis, MDA-MB-231 outcompete MCF-7 cells in physiologic (2 g/L glucose, 7.4 pH) conditions (Figure 2). However, in more acidic conditions (2 g/L glucose, 6.4 pH) the MCF-7 cells appear to persist, suggesting coexistence (Figure 3, Supplementary Figure 1). Furthermore MCF-7 cells persist in either pH condition with high 4 g/L glucose (Figure 4). This raises two questions: 1) Is the success of MDA-MB-231 in most conditions a consequence of higher initial growth rates, increased carrying capacity, or consistently favorable competition coefficients? and, 2) what characteristics of MCF-7 cells allow them to competitively persist at high glucose conditions?

**Figure 2:**
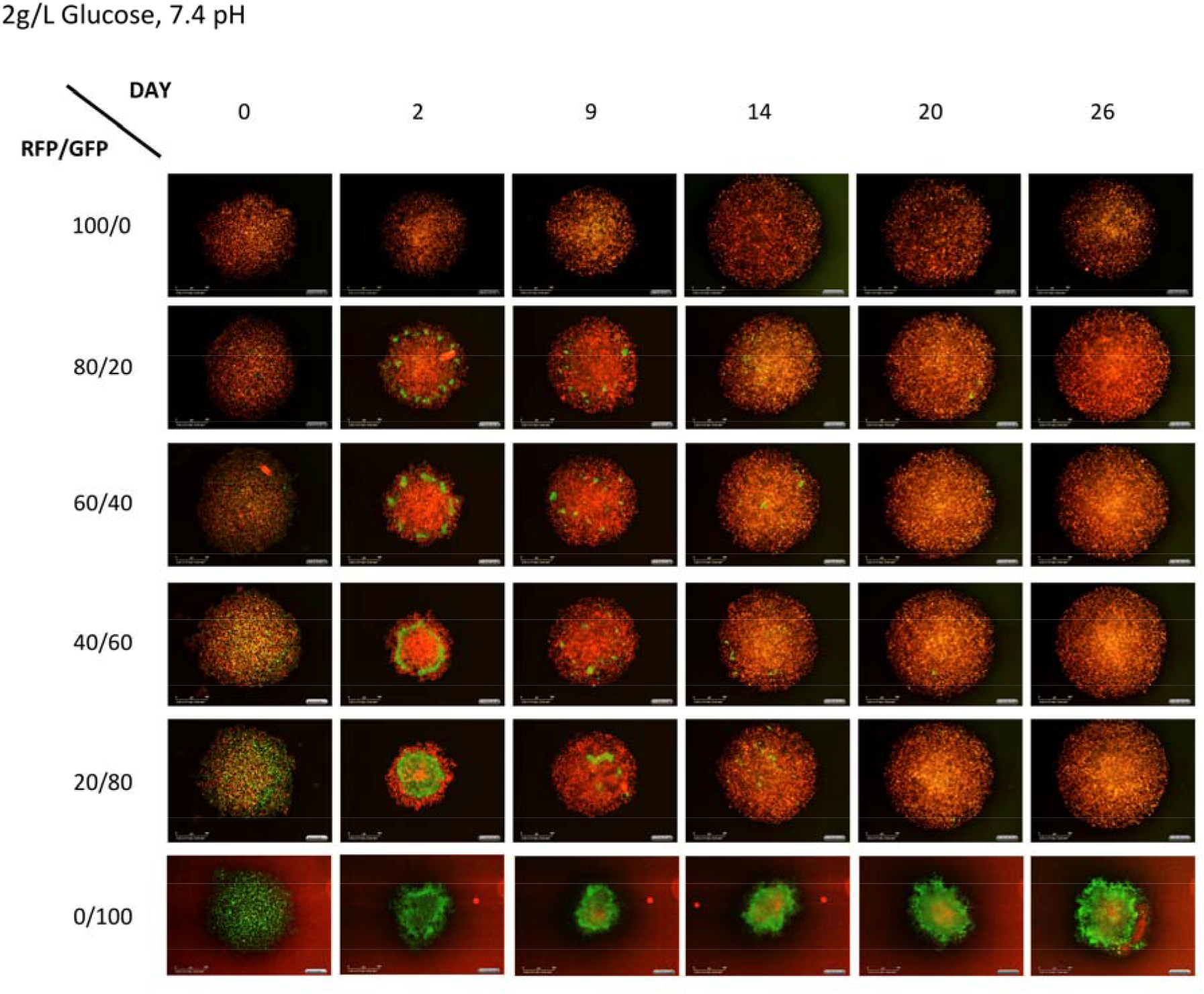
Photographs of spheroids illustrative of the progression and outcome of competition in normal pH. This series shows the progression of spheroids grown physiological conditions (2 g/L glucose, 7.4 pH). Note that, in this case, the MDA-MB-231 cells appear to take over the spheroids by day 20.

**Figure 3:**
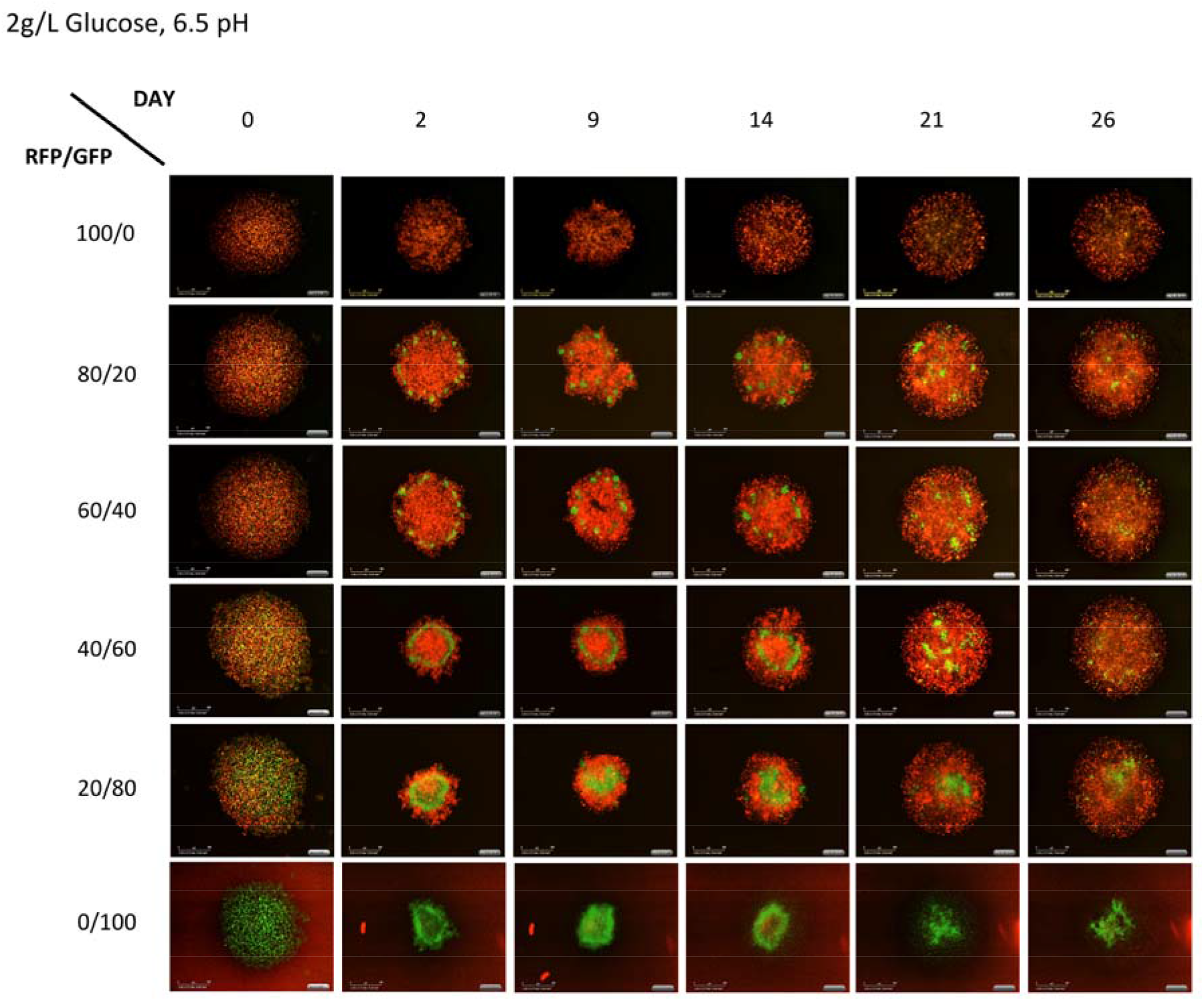
Photographs of spheroids illustrative of the progression and outcome of competition in acidic pH. This series shows the progression of spheroids grown in acidic conditions (6.5 pH) with 2 g/L glucose. Note that, in this case, the MCF-7 cells appear to persist to the end of the experiment.

**Figure 4:**
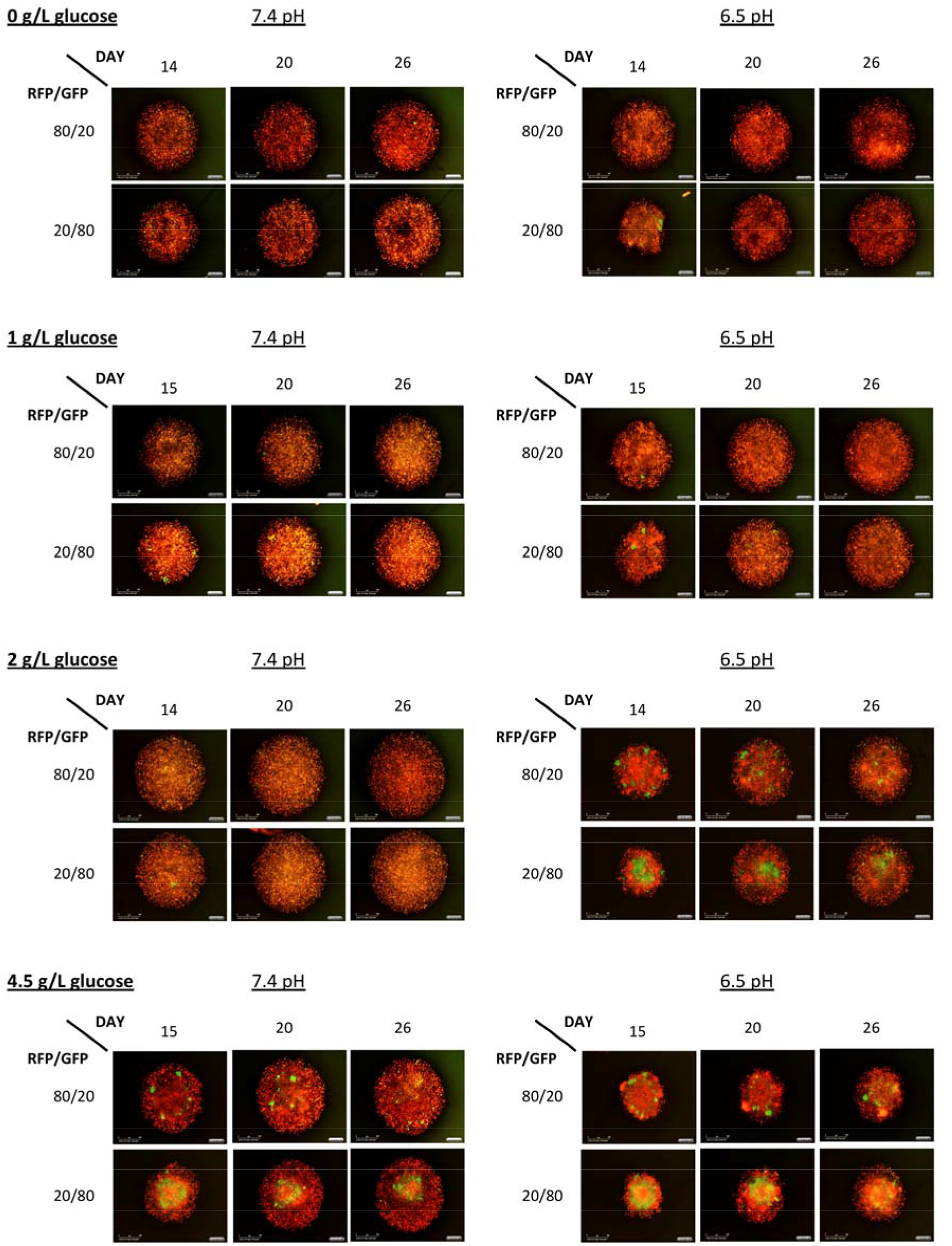
Three time points spanning from the middle to the end for all culture conditions.

#### Analysis of Growth Rate

The growth rate of MCF-7 and MDA-MB-231 cells was quantified by measuring the instantaneous exponential growth rate (in units of per day) for each monoculture during the first 72 hours. In general, MCF-7 cells had equal or higher growth rates under all growth conditions when compared to MDA-MB-231 cells. MCF-7 cell growth rate was reduced when grown in low pH 6.5 (Figure 5 a-c) environments compared to physiologic pH 7.4 (Figure 5 d-f). Not surprisingly, in the harshest environment of low pH 6.5, no glucose, and no glutamine, MCF-7 cells had the lowest, even negative, growth rate (Figure 5c). Interestingly, the absence of glutamine (Figure5 c and f) reduced the difference in growth rates between the two cell types, regardless of pH or glucose concentration, consistent with glutamine’s role as a substrate for respiration, which is more pronounced in MCF-7 cells.

**Figure 5:**
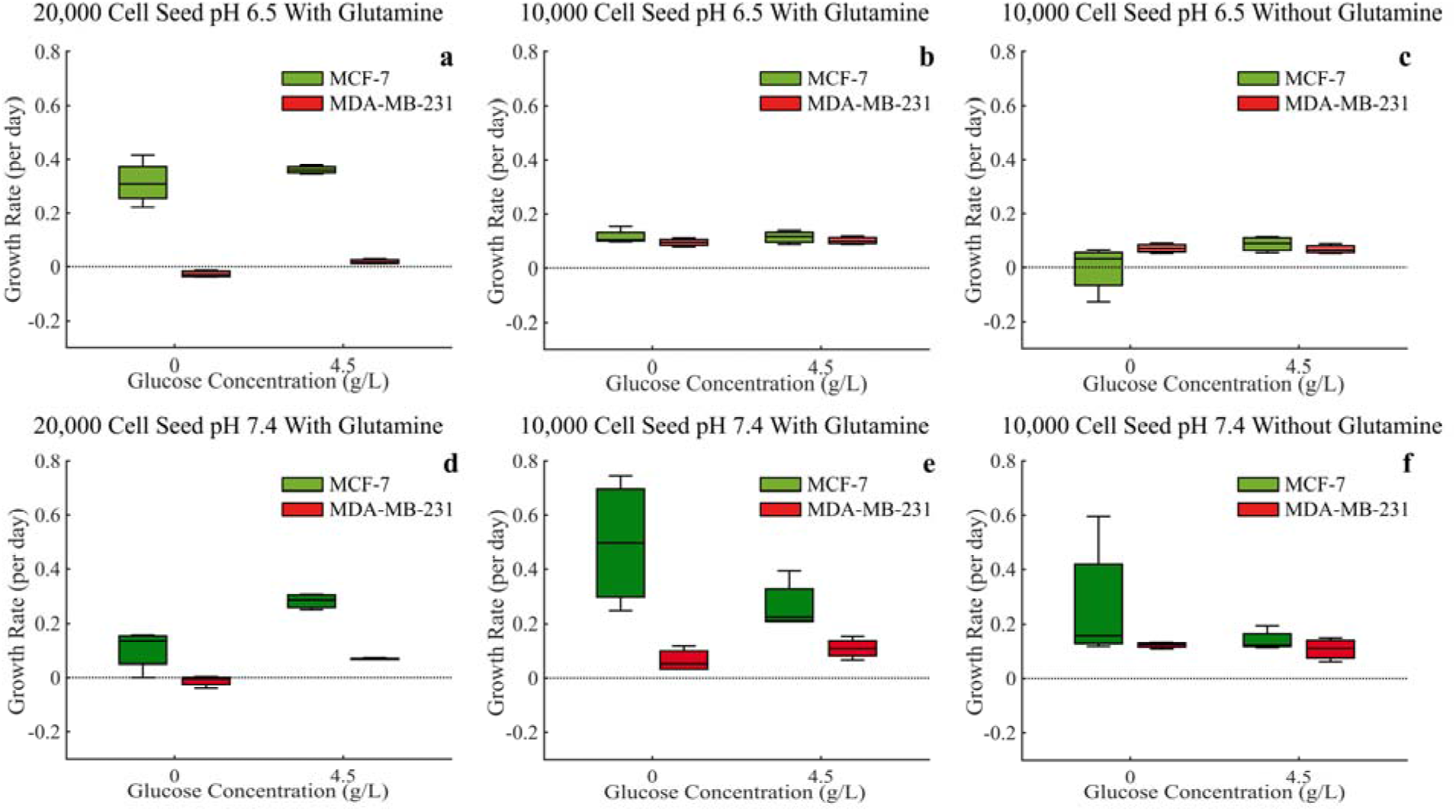
Growth rates of both MCF-7 and MDA-MB-231 cells as measured by the instantaneous exponential growth rate for each monoculture in the first 72 hours under all experimental conditions. ANOVA analysis is provided in Supplemental tables 1-3.

While the growth rates of MDA-MB-231 cells during these early time points were generally unaffected by changes in environmental conditions, the initial plating density had a significant effect. The higher (20,000 initial calls) plating density (Figure 5 a and d) resulted in slower, even negative, growth rates when compared to the lower (10,000 initial cells) plating density. This suggests that these plating densities may be approaching MDA-MB-231’s carrying capacities.

**Table 1:**
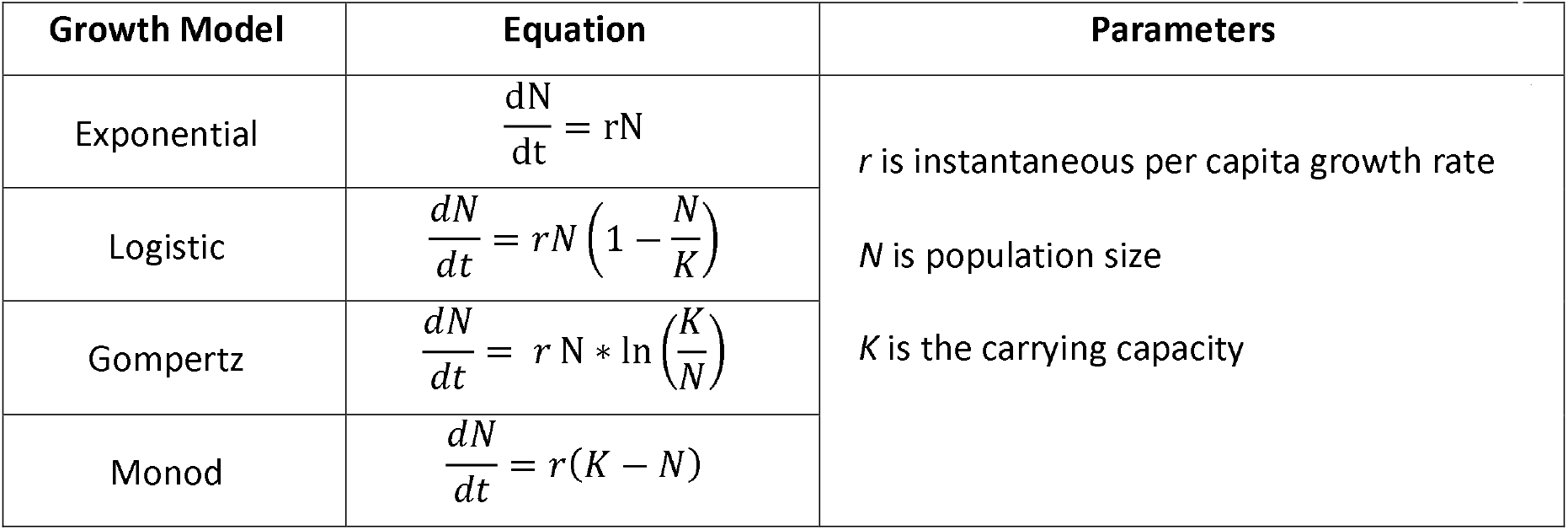
The four models considered for estimating the mode of growth of the cancer cells based on data from monoculture spheroids seeded at a low density of 10,000 cells.

#### Analysis of Carrying Capacity

Due to some differences in the RFU intensity between the GFP and RFP used for the MCF-7 and MDA-MB-231 cells respectively, a direct comparison of carrying capacity using these RFU measurements cannot be made. In this way we split the results in to two figures (Figure 6 and 7). We see RFU’s higher than 60 in the MCF-7 data while the maximum RFU for MDA-MB-231 data is never greater than 20 (please note the varying y-axis limits in the corresponding figures). Even with this difference in RFU intensity, it is not unreasonable to assume, due to the large difference, the MCF-7 cells do indeed have a higher carrying capacity across all experimental conditions.

**Figure 6:**
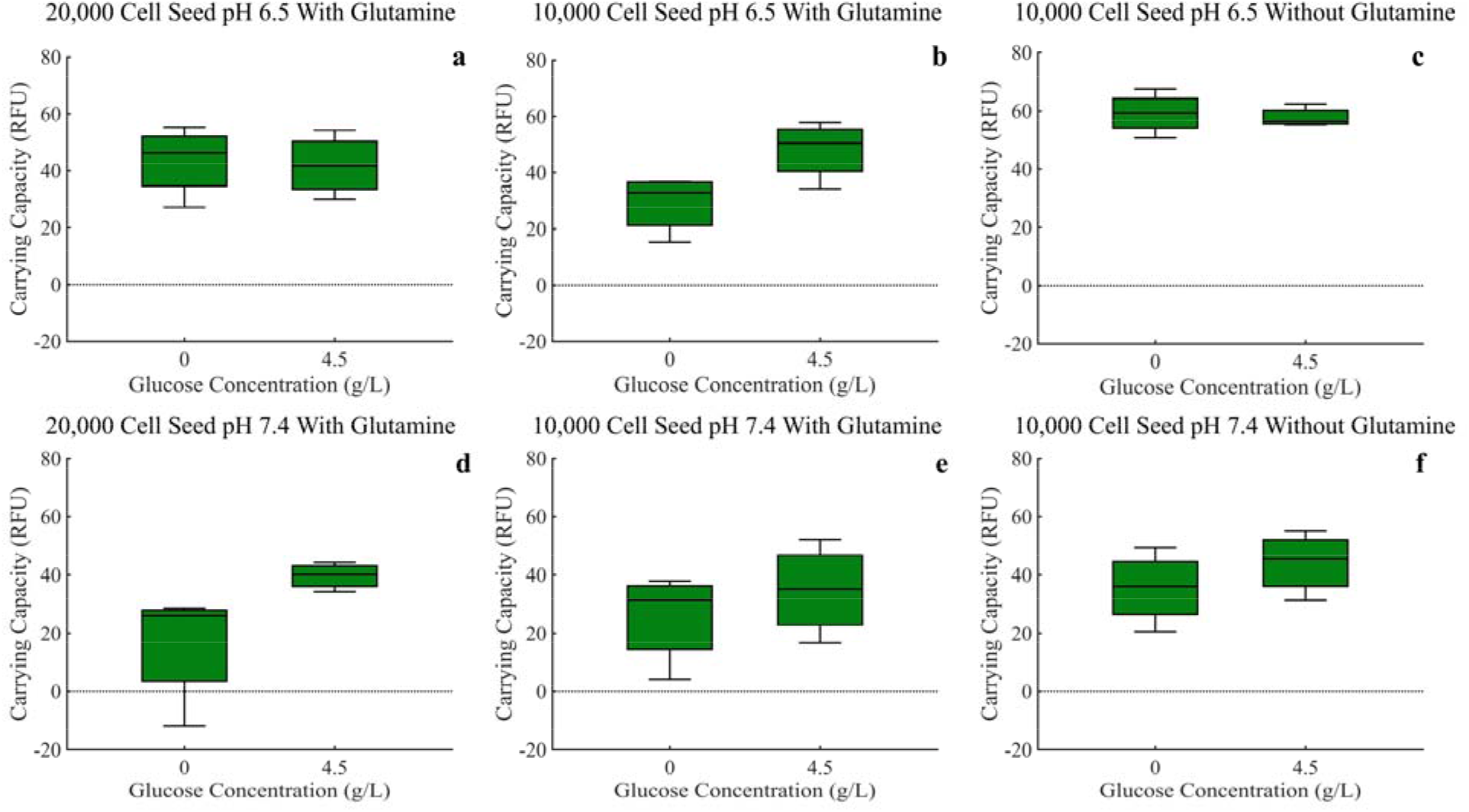
Carrying capacities of MCF-7 cells under all experimental conditions, as measured by the mean fluorescence of the final ten time points. ANOVA analysis is provided in Supplemental tables 4 and 5.

**Figure 7:**
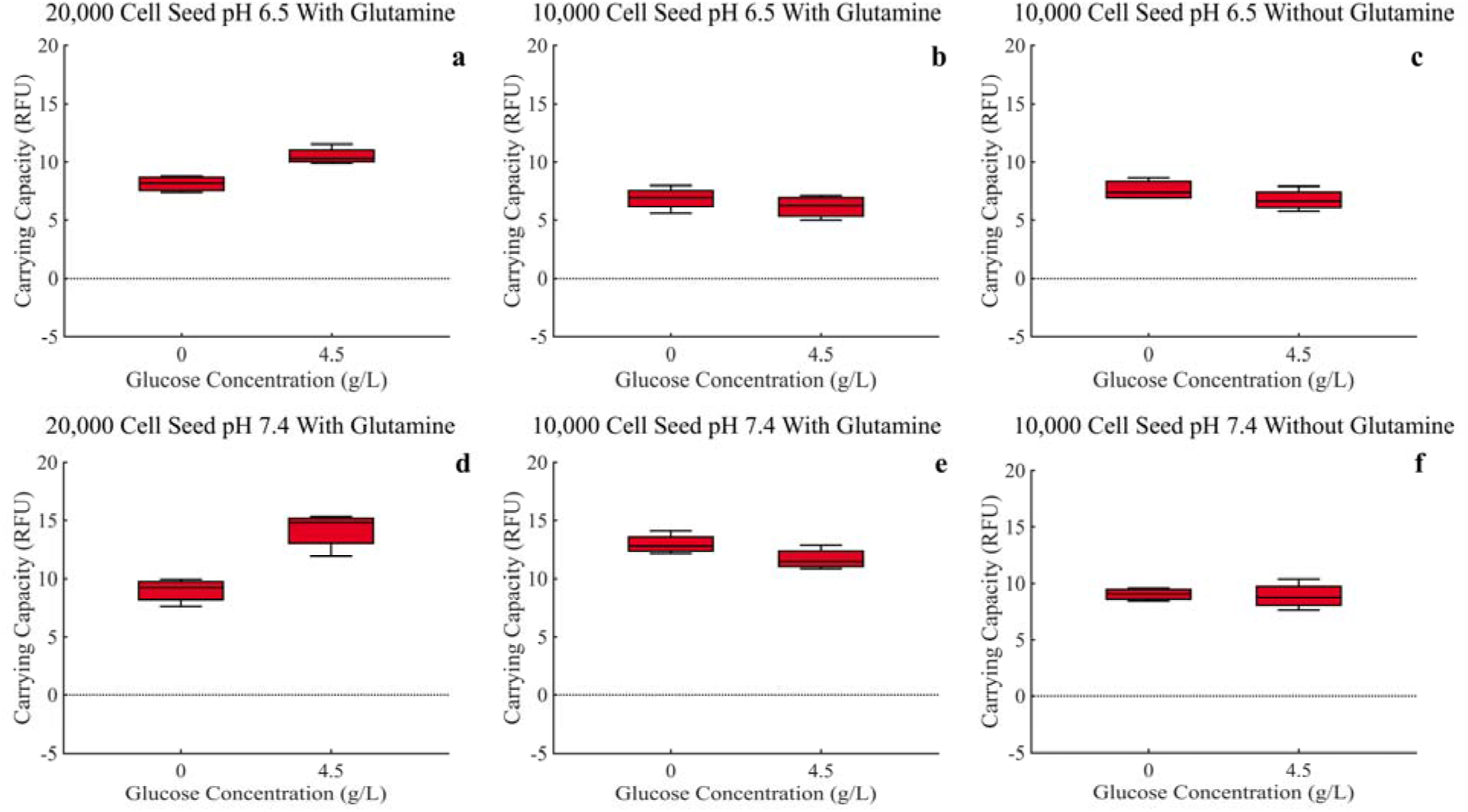
Carrying capacities of MDA-MB-231 cells under all experimental conditions, as measured by the mean fluorescence of the final ten time points. ANOVA analysis is provided in Supplemental tables 4 and 5.

For MCF-7 cells, an increase in glucose resulted in an increase in carrying capacity across experimental conditions, as would be expected (Figure 6 a-f). Surprisingly, the MCF-7 cells experience a decrease in carrying capacity under normal pH conditions when compared to acidic pH 6.5 conditions suggesting an advantage in acidic conditions.

For the MDA-MB-231 cells, after a high seeding density of 20,000 cells, an increase in glucose resulted in an increase in carrying capacity, just like the MCF-7 cells (Figure 7 a and d). For the low seeding density of 10,000 cells, the opposite occurred (Figure 7 b, c, e, and f), though not as dramatically. Furthermore, the MDA-MB-231 cells experience an increase in carrying capacity in normal pH when compared to acidic pH 6.5 conditions suggesting an advantage in normal pH conditions.

#### Analysis of Initial Seeding Frequencies

For the analysis of frequency dependent interactions of the two cells lines we analyzed the instantaneous growth rates (in units of per day) during the first 72 hours in the mixed spheroids (Figs. 8 & 9). The figures are split between low glucose conditions of 0 g/L (Fig. 8) and high glucose conditions of 4.5 g/L (Fig. 9). Interestingly, under both glucose environments, MDA-MB-231 cells have their maximum growth rates when there are only a small initial number of MCF-7 cells in the spheroid. This suggests some benefit to the MDA-MB-231 cells from the presence of MCF-7 cells, or less inhibition of growth rates from MCF-7 cells. Furthermore, we see that the MCF-7 cells generally show a maximum growth rate when the seeding frequencies are 40%/60% or 60%/40% (Figure 8 and 9 b, c, e, and f), again suggesting possible benefits to having both cell types in the spheroid.

**Figure 8:**
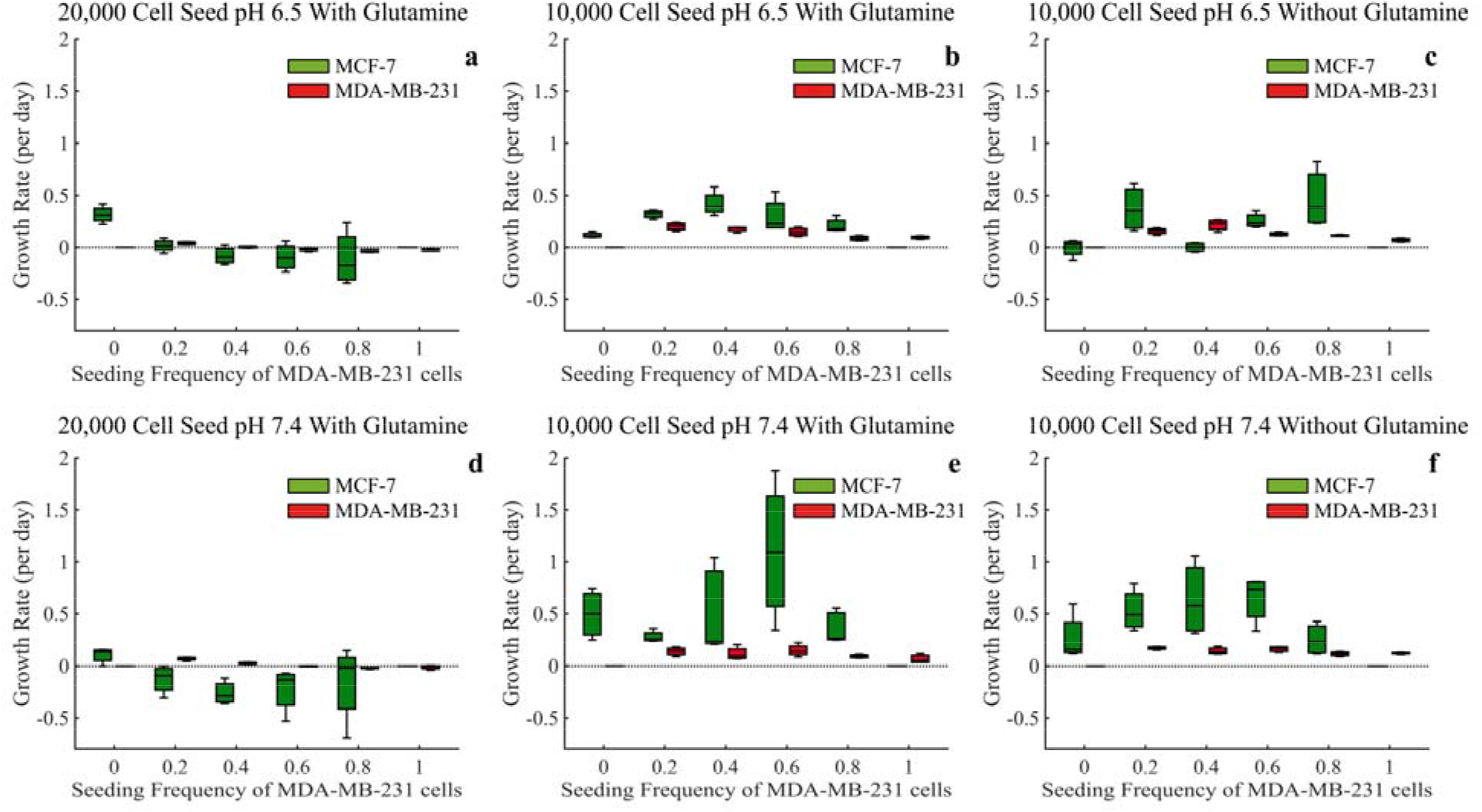
Based on the first 72 hours, estimates of growth rates for mixed population tumor spheroids under glucose starved conditions of 0 g/L. ANOVA analysis is provided in Supplemental tables 1-3.

**Figure 9:**
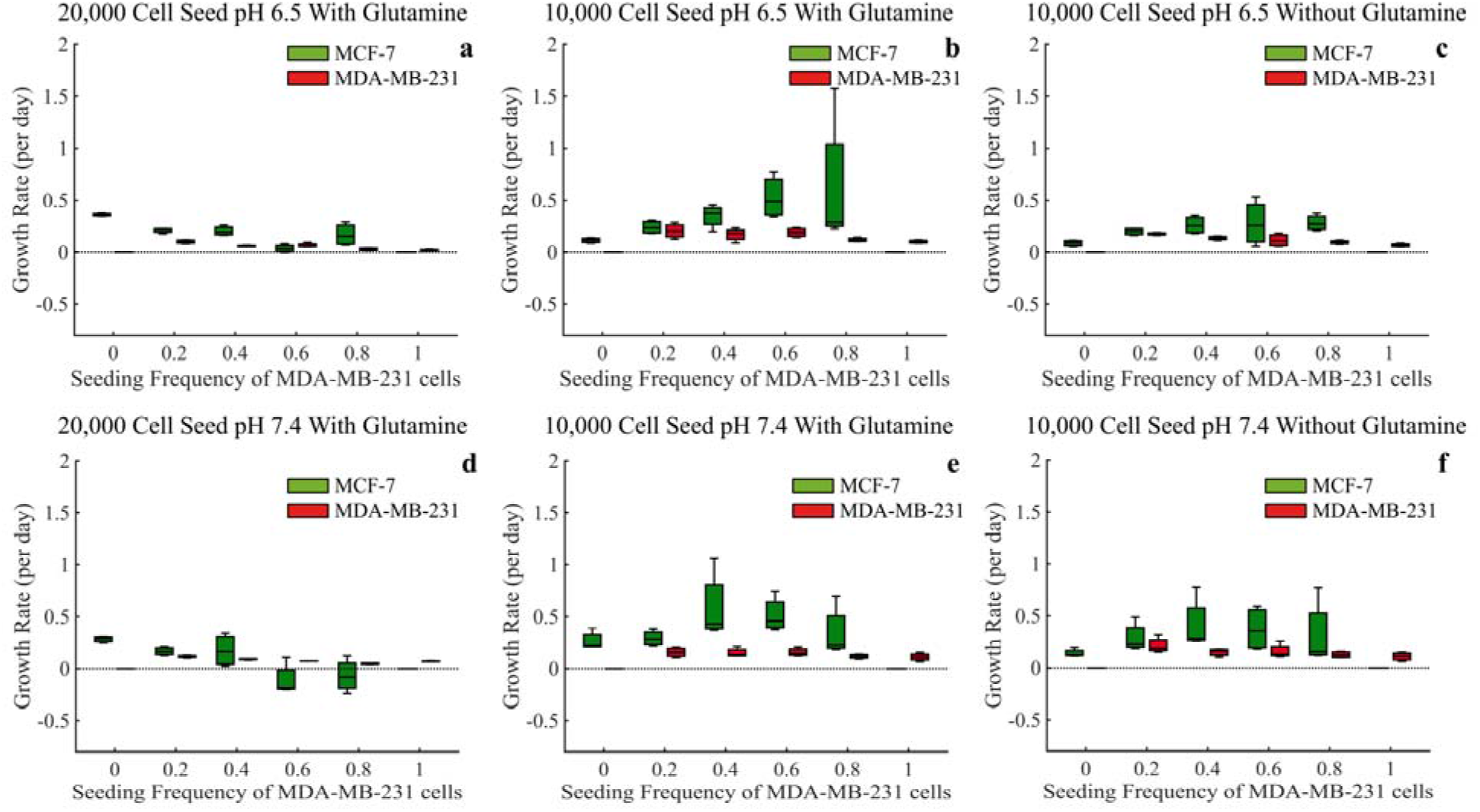
Based on the first 72 hours, estimates of growth rates for mixed population tumor spheroids under glucose rich conditions of 4.5 g/L. ANOVA analysis is provided in Supplemental tables 1-3.

In low glucose and at the high seeding density of 20,000 cells, the MCF-7 cells experience a significant decrease in growth rate, even negative growth rates, as the seeding frequency of MDA-MB-231 cells increase. This points to the MDA-MB-231 cells potentially approaching the carrying capacity provided within these nutrient starved conditions, essentially fully depleting all the resources for other cell types. When the glucose is increased, these MCF-7 growth rates increase, suggesting there are now sufficient nutrients for both cell types (Figs. 8 and 9 a and d). Upon seeding it appears that both cell types generally have higher initial growth rates when the starting frequencies of MDA-MB-231 are low.

#### The Logistic Model Best Describes Individual Cell Growth

Frequency-dependent effects (here measured as competition coefficients) can only be quantified by modeling the growth of multiple interacting types or species. In order to obtain an accurate estimate of these, the mode of growth must first be determined. In this way, we use the monoculture spheroids of either MCF-7 or MDA-MB-231 alone as a data set to first understand the growth dynamics without the added complexity of the mixed cultured spheroids. Although several growth models have been proposed for spheroids, tumors, and organoids, no specific consensus has been reached (34, 35). Because of this, we agnostically compared four population growth models: exponential, logistic, Gompertz, and Monod-like (Table 1).

The exponential growth model is commonly used in bacterial studies and assumes unlimited resources. Logistic population growth is the simplest constrained growth model and assumes that per capita population growth rate declines linearly with population size or density (36). Although it gives a similar shape, Gompertz growth is exponentially constrained by increasing population. Several authors and works have suggested that the Gompertz equation provides a better fit to the growth of tumors *in vivo* (37). The Monod-like equation imagines resource matching where per capita growth rates are proportional to the amount of resources supplied per individual. It provides a good fit to the population growth curves of bacteria and other single cell organisms grown in chemostats (38). Here, we consider all four models when estimating the DI parameter of intrinsic growth rate (r) and the latter three models for estimating the DD parameter of carrying capacity (K).

Using constrained parameter optimization, all four models were fit to the mono-culture spheroid growth of MCF-7 or MDA-MB-231 cells under the eight environmental conditions (Supplementary Figure 1). Due to the lower number of time points in the experiments with a high seeding density of 20,000 cells, these experiments were not used. Quality of fit for each model was evaluated using adjusted R2, Root Mean Square Error (RMSE), and Akaike Information Criterion (AIC) (Table 2, Supplementary Table 6).

**Table 2:**
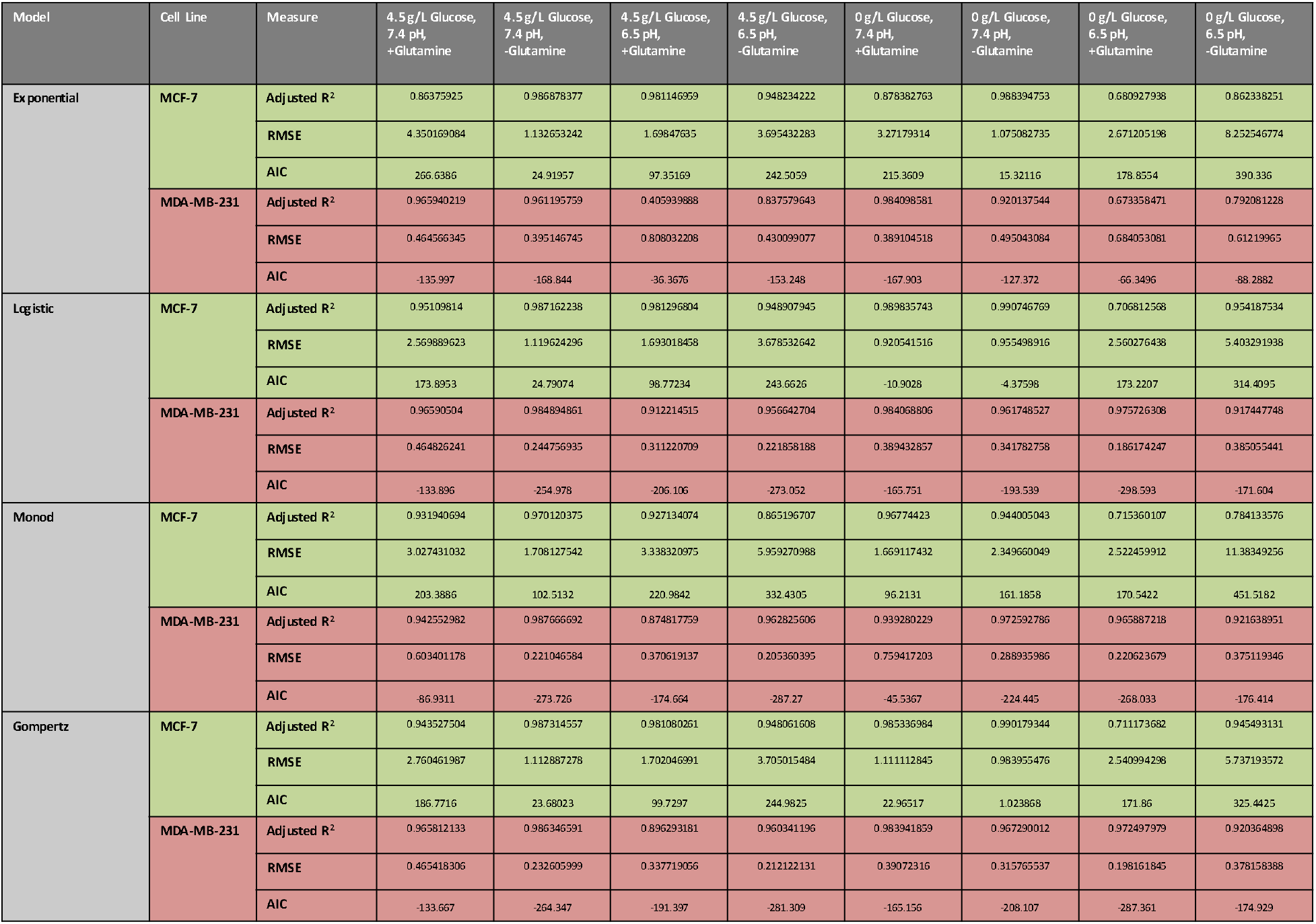
Average adjusted R^2^, RMSE, and AIC when fitting the four growth models to monoculture spheroids.

Two-way ANOVAs (one for each cell type with pH and nutrient levels as independent variables) showed that adjusted R^2^’s were similar for the three density-dependent models and significantly lower for the exponential model (Supplementary Table 6). Conversely, the RMSE indicated the best fit for exponential growth (lowest value), lowest quality fit for the Monod-like equation, and similar, intermediate values for the logistic and Gompertz growth models. AIC analysis suggests logistic fits to be the best fit in the majority of scenarios.

While any of these four models could be used to model this system, this analysis suggests either an exponential or logistic model provides the best fit to the data. The principal difference between these two models is that logistic growth includes density-dependence and the growth curve approaches a carrying capacity, whereas exponential growth ultimately leads to extinction (if negative growth rates) or infinitely large population sizes (if positive growth rates). In this way, we evaluated whether the spheroids showed evidence of approaching or reaching a carrying capacity. To do this, we examined the slope of the final ten time points for both the red and green fluorescent values (RFU’s) for all experimental conditions (Supplementary table 7). As most mono-culture spheroids showed declining slopes, or slopes near to zero, we favor the logistic model for this study.

#### Parameter Estimation for Lotka-Volterra Competition Model

With the logistic equation showing the best fit for the mono-culture experimental data, we expanded to the Lotka-Volterra competition equations to model the mixed population tumor spheroids. The Lotka-Volterra equations represent a two species extension of logistic growth with the addition of competition coefficients.

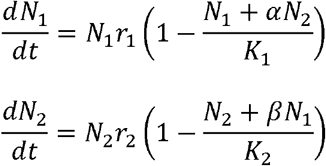

where *N_i_* are population sizes, *r_i_* are intrinsic growth rates and *K_i_* are carrying capacities. We let species 1 and 2 be MCF-7 and MDA-MB-231, respectively. The competition coefficients (α and β) scale the effects of inter-cell-type competition where α is the effect of MDA-MB-231 on the growth rate of MCF-7 in units of MCF-7, and vice-versa for β. If α (or β) is close to zero, then there is no inter-cell-type suppression; if α or β) is close to 1 then intra-cell-type competition and inter-cell-type competition are comparable; and if α (or β) is >1 then inter-cell-type competition is severe.

The values of the carrying capacity of MCF-7 cells *K_MCF_*, the carrying capacity of MDA-MB-231 cells *K_MDA_*, and the competition coefficients, α and β, predict the outcome of inter-specific competition. MCF-7 cells are expected to outcompete MDA-MB-231 cells if *K_MCF_* > *K_MDA_* / β and *K_MCF_* / α > *K_MDA_*. Similarly, MDA-MB-231 cells are expected to outcompete MCF-7 cells if *K_MDA_* / β > *K_MCF_* and *K_MDA_* > *K_MCF_* / α. Either MCF-7 or MDA-MB-231 will out-compete the other depending upon initial conditions (alternate stable states) when *K_MCF_* > *K_MDA_* / β and *K_MDA_* > *K_MCF_* / α. The stable coexistence of MCF-7 and MDA-MB-231 cells is expected when *K_MDA_* / β > *K_MCF_* and *K_MCF_* / α > *K_MDA_*.

To estimate the values for *K_MCF_*, *K_MDA_*, α, and β, we fit the co-culture data with low seeding density to the two-species Lotka-Volterra equation creating different fits for each combination of experimental conditions. It is important to note here that the Lotka-Volterra competition equations are not spatially explicit. From Figures 2-4, it can be seen that the cells are segregated within the tumor spheroid. In our application, the Lotka-Volterra competition equations simply provide a first-order linear approximation of between cell-type competition, not a mechanistic model of the consumer-resource dynamics or the spatial dynamics that likely occur in the spheroids.

Using nonlinear constrained optimization, we varied the values for instantaneous growth rates, carrying capacities, and competition coefficients α and β to minimize the RMSE between the Lotka-Volterra model and the experimental data (Figure 10). In addition to estimating the competition coefficients, this uses the mixed culture data to re-estimate the growth rates and carrying capacities previously estimated from the monoculture spheroids as growth rate and carrying capacity depend on initial seeding density (Figs 8-9).

**Figure 10:**
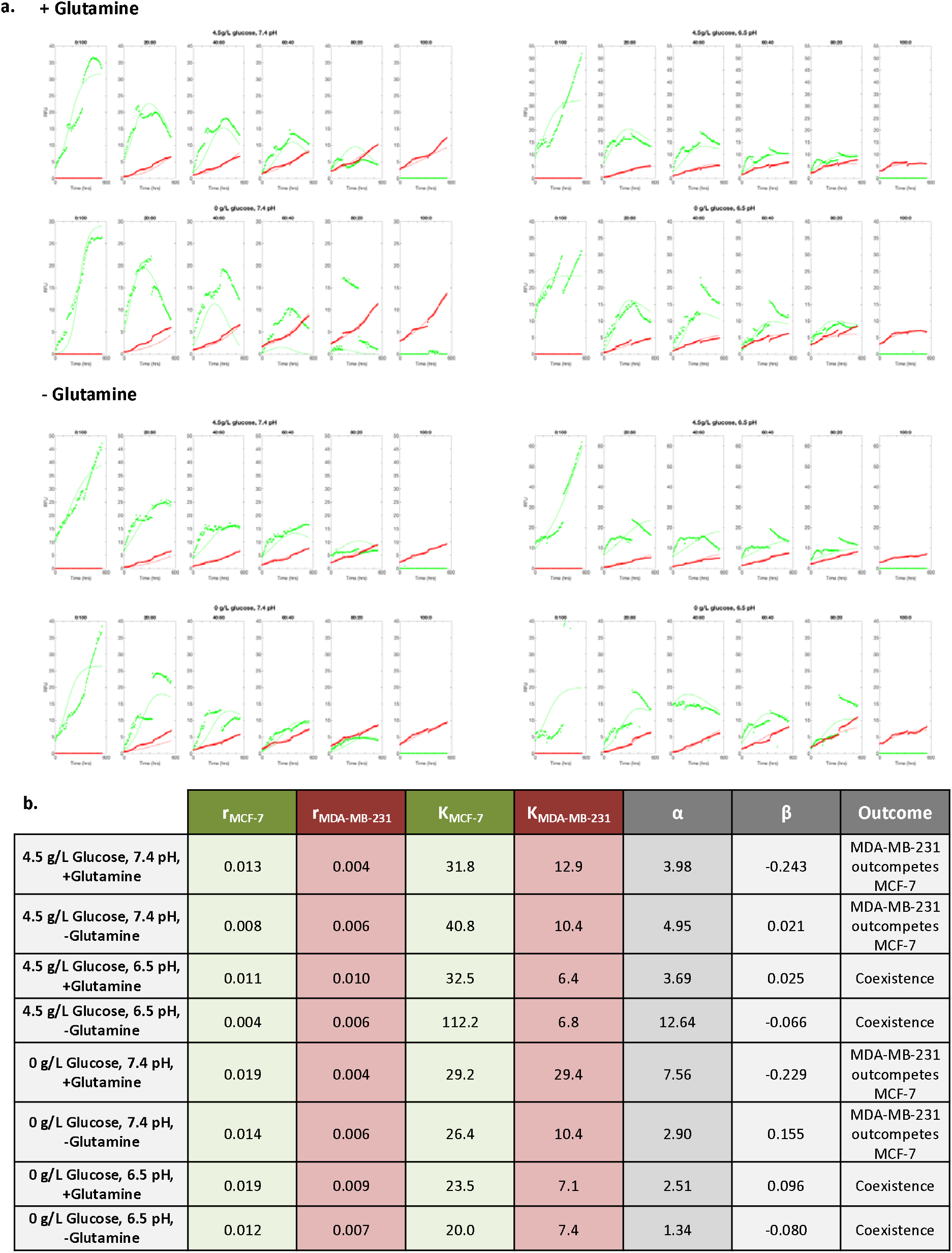
**A.** Estimation of growth rates, carrying capacities, and competition coefficients using nonlinear constrained optimization fitting to the Lotka-Volterra competition growth curves to the low (10,000 cell) seeding density experiments. The average of the four replicates for each experimental condition are shown along with the optimized fit to the Lotka-Volterra model. **B.** Optimized parameters for growth rates, carrying capacities, and competition coefficients are shown for each experimental condition. The predicted outcome of competition predicted by these parameters is given in the last column.

As in the monoculture parameter estimation, MCF-7 had higher intrinsic growth rates and carrying capacities than MDA-MB-231 under all culture conditions. For all culture conditions, α > 1 (as high as 12.6) and β ≈ 0. This indicates that the MDA-MB-231 cells had large competitive effects on MCF-7 growth, and that MCF-7 cells had virtually no inhibitory effects on the growth of MDA-MB-231 cells. It appears that this frequency dependent (FD) effect provides a competitive advantage for the MDA-MB-231 cells that allows them to remain in the tumor spheroids despite MCF-7’s higher intrinsic growth rates (DI) and carrying capacities (DD).

The greatest frequency dependent effect of MDA-MB-231 on MCF-7 (highest α) occurred under conditions of 4.5g/L glucose, 6.5pH, and no glutamine. The greatest effect of MCF-7 cells on MDA-MB-231 (highest β) occurred in the absence of glucose and glutamine at neutral pH. This fits our original predictions; however, the effect of β is not strong enough to rescue MCF-7 cells from competitive exclusion. In accordance with the hypothesis that glucose permits MDA-MB-231 cells to be highly competitive, α’s (the effect of MDA-MB-231 on MCF-7) were generally lower under glucose-starved conditions. Further, both cell types appear to experience decreased growth efficiency and competition when glucose/glutamine starved, and under acidic conditions. This is consistent with more general ecological studies showing that competition between species is generally less under harsh physical conditions (39, 40).

Using the optimized parameters values calculated by the constrained optimization, analysis of the Lotka-Volterra equations can predict the outcome of competition. This analysis suggests that under physiologic pH, regardless of glucose or glutamine concentration, MDA-MB-231 cells will outcompete the MCF-7 cells. Furthermore, under acidic pH, regardless of glucose or glutamine concentration, MDA-MB-231 cells and MCF-7 cells will actually coexist. This is observed experimentally in Figure 2 and Figure 3 where acidic pH results in coexistence of the two cell types and normal pH results in no MCF-7 cells remaining at the end of the experiment. While a robust result, this does not accord with our original expectations.

#### In Vivo Exploration of Cell Competition

MDA-MB-231 and MCF-7 cells were grown in the mammary fat pads of 8-10 week old nu/nu female mice. Three tumor models were evaluated: only MCF-7, only MDA-MB-231, and 1:1 mix of both cell types with three replicates for each. Tumor volume was measured weekly. At five weeks, the tumors were harvested and evaluated for ER expression which marks MCF-7 (ER positive) cells providing estimates of the ratio of MDA-MB-231 to MCF-7 cells.

H&E staining indicates that MCF-7 tumors had the greatest viability and lowest necrosis while MDA-MB-231 tumors displayed decreased viability and increased necrosis (Figure 11a). The small differences in necrotic and viable tissue between the MDA-MB-231 and mixed tumors are indicative of the expansive effect that MDA-MB-231 phenotype has on tumor progression (Figure 11c).

**Figure 11:**
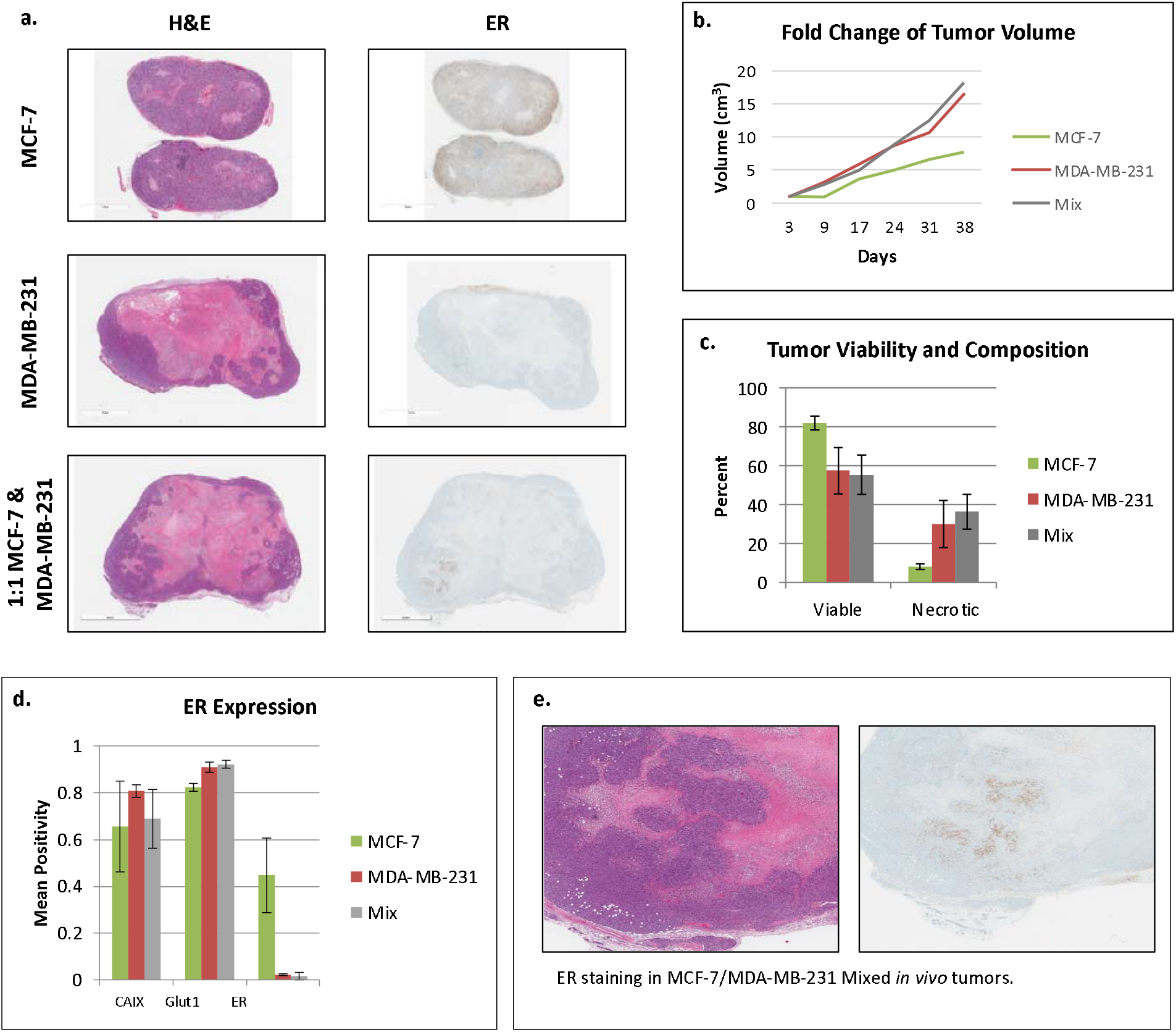
Results of *in vivo* mono- and co-culture tumors. **A.** Histology slides of MCF-7, MDA-MB-231, and 1:1 co-cultured tumors. Slides were stained with H&E and ER immune stain. **B.** Average fold change of tumor volume reflects relative growth. **C.** Tumor viability for all three tumor types. Notice a decrease in viability and increase in necrosis in the MDA-MB-231 and mixed tumors. **D.** Measures of CAIX, Glut1, and ER biomarker staining for each tumor type. **E.** The presence of small clumps of ER staining in mixed tumors reflect the clumping of MCF-7 cells in the progression of the mixed tumor spheroids, suggesting the coexistence of these two phenotypes in certain tumor microenvironments.

Changes in tumor size over five weeks indicated that MCF-7 cell tumors grew more slowly than MDA-MB-231 tumors and mixed tumors (Figure 11b). This is an interesting result as the in-vitro analysis would have predicted that the MCF-7 tumors would grow the fastest due to both higher growth rate and higher carrying capacity. Only in the harshest in-vitro condition with 0 g/L glucose, acidic pH 6.5, and no glutamine did MCF-7 cells grow more slowly than MDA-MB-231. This potentially suggests that mammary fat pads may exhibit a harsh environment for MCF-7 cells.

Of interest, the mixed tumors in-vivo grew significantly faster than the MCF-7 tumors, and potentially faster than the MDA-MB-231 tumors alone beyond 24 days. This reiterates the in-vitro experiments in Figures 8 and 9 where both cell types gain some growth advantage when grown together.

Biomarker staining indicated the presence of microenvironment regulating membrane transporters. Variation in CAIX expression between the three tumor types suggests little difference in active pH regulation. Glut1 expression is higher in MDA-MB-231 containing tumors, reflecting the increased capacity for glucose uptake by these cells.

ER staining was also used to identify the final frequency of MDA-MB-231 and MCF-7 cells. The amount of ER staining in the mixed tumors (mean of 0.0207) was comparable to the amount found in the MDA-MB-231 only tumor (mean of 0.0227), indicating significant loss of MCF-7 clones at the experiment’s end point. This reiterates the results from the in-vitro spheroid experiments demonstrating that MDA-MB-231 cells have a competitive advantage. Interestingly, in one of the mixed tumors (Figure 11e), we note the presence of a cluster of ER positive cells concentrated in an area of interior viable cells. From in-vitro and theoretical analysis, we predict that coexistence of these two cells lines is possible under acidic conditions. In the presumably acidic interior of the in-vivo tumor, while the density of the MCF-7 cells is low, coexistence may be occurring. This clumping MCF-7 cells *in vivo* closely matches the mixed culture *in vitro* spheroids. The spatial distributions of the two cell types may be an important component for coexistence.

## DISCUSSION

To understand competition among cancer cell phenotypes within a tumor, we hypothesized that direct competition experiments are critical for understanding factors that may promote or inhibit coexistence. Similar to the classic experiments by Gause, we conducted competition experiments between two well-established breast cancer cell lines that exhibit classic phenotypic and in-vivo growth patterns of the 1) “Pioneer” MDA-MB-231 cells, which are highly invasive, non-angiogenic, and glycolytic and the 2) “Engineer” MCF-7 cells which are angiogenic, non-invasive, estrogen dependent and have near normal glucose metabolism.

While here we focused specifically on nutrition and pH as factors that affect cellular competition, there are a number of other factors that undoubtedly play a role. For example, hormonally driven cell lines might elucidate the role of systemic and intratumoral hormone levels in competition experiments such as T47D (ER+, PgR+, HER2−) as another “engineer” instead of MCF7. Or interestingly, one could use the mouse breast cancer cell line MMTV-PyMY (ER−, PgR− Her2+/−) as a “pioneer” instead of MDA-MB-231 cells to elucidate the protein-protein interaction between human and mouse cells. In actuality, any number of cancer cell lines would be good candidates for competition assays covering additional factors that might affect cellular growth and viability.

Here, our first experiments investigated monocultures in various nutrition and pH conditions likely to be present in some regions of an *in vivo* tumor. In our experimental spheroid model, the growth rate (*r*) and carrying capacity (*K*) for MCF-7 cells in monoculture were the same or greater than those of MDA-MB-231 cells under all conditions. We expected these advantages would allow the MCF-7 cells to dominate the MDA-MB-231 cells in co-culture, but surprisingly, MDA-MB-231 cells dominated in all experimental conditions. This counterintuitive result was also found in-vivo, where MCF-7 tumors generally grew slower than MDA-MB-231 tumors. This suggested that the density dependent and density independent interactions can not fully explain cell competition.

In this way, the frequency dependent interactions become paramount. We found that magnitude of this frequency dependent competitive effect of one cell type on the other is highly dependent on culture conditions. In spheroids, MDA-MB-231 cells displayed their greatest competitive effect in high glucose and low pH culture conditions while MCF-7 cells, although overall less competitive, showed their greatest competitive effect in low glucose and neutral pH culture conditions. This aligns with the “pioneer” and “engineer” phenotypic patterns and suggests that tumor heterogeneity is driven not only by clonal growth efficiency but also by the landscape of the tumor microenvironment. The mechanisms of this competition is unknown, but there are a number factors that could play a role beyond those tested here such as cytokine production, growth and inhibitory signal dynamics, and direct contact-dependent signals to name a few.

In general, the outcome of competition appeared to be pH specific with MDA-MB-231 cells dominating in normal pH conditions and MCF-7 and MDA-MB-231 cells coexisting in acidic conditions both in-vitro and in-vivo. This is evident not only in the overall growth dynamics of competition between MCF-7 and MDA-MB-231 cells, but also by the spatial patterns observed both *in vitro* and *in vivo*. The emergence of such spatial patterns suggests that some aspects of intratumoral spatial distribution are the result of cellular phenotypic properties. In the co-culture spheroids and in the example shown in-vivo, MCF-7 cells form a ring or cluster of highly clumped cells away from the spheroids edge. The MDA-MB-231 cells, on the other hand, formed the spheroid or tumor edge by extending their range outward beyond that of the MCF-7 cells. This same pattern, where cells on the periphery of a tumor leave an open niche for another cell type in the interior, has been shown experimentally and also analyzed using Lotka-Volterra competition equations in the *Drosophila* imaginal disc (41–43).

The competitive superiority of MDA-MB-231 over MCF-7 from our in vitro and in vivo experiments suggests new hypotheses. Several eco-evolutionary mechanisms may be responsible. In the competition experiments, the cancer cells experienced pulsed resource renewal every 3-4 days. Thus, over a four day period the culture media likely went from nutrient rich to poor, while metabolite concentrations, including acid, increased (44). In both natural and malignant ecosystems, oscillating resources and metabolites favor traits of 1) “priority”, 2) speed, and 3) efficiency (45–47).

Here, *priority* is defined by the ability to localize, move to, and invade areas of high resource availability. The higher motility and invasiveness of the MDA-MB-231 allowed them to occupy the outer margins of the spheroid where nutrient concentrations would be highest (48). *Speed* is defined by the rate at which an organism can forage. A high speed allows an organism to pre-empt other organisms via interference with uptake by competitors (49). The high glycolytic capacity of the MDA-MB-231 permits them to acquire glucose faster than their MCF-7 competitors on a per cell basis. Furthermore, MDA-MB-231 do not adhere to each other as the MCF-7 cells did. This gives the MDA-MB-231 a higher surface area for nutrient uptake and less cell-to-cell interference in nutrient uptake. Finally, the motility and speed of nutrient uptake may both contribute to rapid but inefficient foraging. High speed and efficiency benefit the cell both through acquisition of resources and by making them unavailable to its competitors.

However, high motility and speed of nutrient uptake often result in rapid but inefficient foraging. The lower intrinsic growth rates and carrying capacities of MDA-MB-231 are consistent with this. *Efficiency* is the capacity of an organism to cheaply and profitably exploit resources when they are scarce. The MCF-7 may be slower, more efficient foragers thus depleting nutrients following the pulse more slowly but in a manner that allowed for greater per cell survival and proliferation, thus explaining their higher growth rates and carrying capacity when grown in mono-culture (50, 51).

The outcome of competition between the two populations were almost entirely frequency dependent as reflected in the highly asymmetric and strong competitive effect of MDA-MB-231 cells on MCF-7 cells. Yet, we saw an important role for MCF-7’s higher carrying capacity (density-dependent effects) in promoting coexistence under some circumstances, and the differences in intrinsic growth rates (density-independent effects) likely influenced the transient dynamics within the mixed cultures.

## CONCLUSIONS

Mixed-culture competition experiments in spheroids can successfully track competitive outcomes between cell lines of interest (52–54). Because alterations in glucose and pH can significantly affect the competitive ability of different cell phenotypes, manipulation of the microenvironment via therapies could directly influence intra-tumoral evolution and cell-type composition (55–57). Such experiments in spheroids may be valuable for quantifying the costs of resistance. Experiments could consider competing a resistant cell line against its sensitive parental cell line under conditions of no therapy (sensitive would be expected to outcompete the resistant cell line) and under conditions of therapy (resistant should outcompete the sensitive cell line) (58). As accomplished here, one could determine differences in growth rates, carrying capacities and competition coefficients. These metrics would be valuable for informing adaptive therapy trials that modulate therapy so as to maintain a population of sensitive cells as a means for competitively suppressing the resistant cells (59–61).

Mixed culture competition experiments demonstrate how both the densities and the frequencies of cell types influence the growth rates and population dynamics of cancer cells. Measurements of resource availability and carrying capacities determine limits to growth. The extent to which inter-specific competition coefficients differ significantly from 1, reveal the extent to which cancer cells engage in eco-evolutionary games where the success of a given cancer cell depends both on its phenotype but the phenotypes of the other cancer cells. Such understanding of how cancer cell types respond to each other and to therapy will be crucial to evolutionarily informed therapies that aim to anticipate and steer the cancer’s eco-evolutionary dynamics.

## METHODS

### Cell Culture and acid adaptation in vitro

MCF-7 and MDA-MB-231 cells were acquired from American Type Culture Collection (ATCC, Manassas, VA, 2007–2010) and were maintained in DMEM-F12, no glutamine, no glucose, no HEPES (Life Technologies) supplemented with 10% FBS (HyClone Laboratories). Cells were tested for mycoplasma contamination and authenticated using short tandem repeat (STR) DNA typing according to ATCC guidelines that is approved by Moffitt Cancer Center. Growth media was supplemented to contain adjusted glucose and glutamine concentrations, depending on desired environmental conditions. Media was also supplemented with 25 mmol/L each of PIPES and HEPES to increase the buffering capacity of the media and the pH adjusted to 7.4 or 6.5 for each spheroid growth condition.

### Transfection (GFP/RFP plasmids)

To establish stable cell lines, the MCF-7 and MDA-MB-231 cells were infected with Plasmids expressing RFP or GFP using Fugene 6 (Promega) at an early passage and were selected using 2 μg/ml puromycin (Sigma).

### 3D spheroid co-culture

Perfecta3 96-Well Hanging Drop Plates or non-adhesive U shape bottom 96 well plates were used to grow the primary spheres containing 10,000 or 20,000 total cells. For each experimental condition, the ratio between MCF-7-GFP and MDA-MB-231-RFP was from 0 to 100 percent as follows: 0/100, 20/80, 40/60, 60/40, 80/20, 100/0. Each ratio had four replicates for each experiment. An Incucyte microscope kept at 37oC and 5% CO2 was used to image the spheroid growth over approximately 30 days. Images and fluorescent intensity were analyzed in Incucyte built-in software and ImageJ (62). Relative fluorescent units (RFUs) were normalized by averaging red/green fluorescence when the corresponding cell type was absent then subtracting this value from other fluorescent values of that cell type.

### Experimental conditions

Two experiments were conducted. The first consisted of spheroids with a starting density of 20,000 cells under conditions of high (4.5g/L), medium (2g/L), intermediate (1g/L) and no glucose fully crossed with physiological (7.4) and low (6.5) pH. Spheroids were allowed to grow for roughly 670 hours with fluorescence measured every 24-72 hours. Relative fluorescent units (RFUs) were used to generate estimates of growth rates and carrying capacities.

Observations from these spheroids prompted us to conduct a second set of experiments consisting of spheroids with a lower starting density (10,000 cells) cultured under the extremes of nutrient concentration (0 g/L and 4.5 g/L glucose) fully crossed with physiological and low pH. Resilience of growth under no glucose prompted a repeat of the experiment 0 g/L and 4.5 g/L glucose fully crossed with physiological and low pH with medium lacking glutamine. To improve accuracy of model-fitting, fluorescent values were collected every 6 or 12 hours.

#### Calculation of growth rate and carrying capacity of mono-cultures

Growth rates were calculated as the log change of the first 72 hours after plating. Carrying capacities were calculated as the mean fluorescence of the final ten time points for each experiment. Since their values were similar regardless of starting frequency, and since they dominated the tumor spheroid in all experiments, we could estimate the carrying capacities of MDA-MB-231 cells using all cultures with a starting frequency of > 20% MDA-MB-231. Due to competitive effects, MCF-7 carrying capacities were estimated using only the 100% MCF-7 spheroids.

### Testing among growth models

Nonlinear constrained optimization was used for parameter estimation of r and K to identify the best fit of the monoculture experiments. The objective function minimized the RMSE between the theoretical curves for exponential, Monod-like, logistic, and Gompertz growth models to the growth dynamics of each monoculture spheroids growth dynamics.

### Competition Coefficient Estimation

Nonlinear constrained optimization was used for parameter estimation of the two-species competition Lotka-Volterra equations. The objective function minimized the cumulative RMSE of all initial seeding densities in order to obtain the parameters that could best explain all experimental initial conditions. For these estimations we allowed the optimization technique to optimize all four parameters: r, K, α, and β. While we could have fixed the growth rates and carrying capacities using the values found during the growth model analysis, we instead allowed them to also vary due to the experimental evidence showing that initial seeding frequency can significantly change the growth rates and carrying capacities of these cells.

### In Vivo Tumor Culture

MCF7 and MDA-MB-231 breast cancer cell lines were purchased from ATCC. Both cells were authenticated by short tandem repeat analysis and tested for mycoplasma. Xenograft studies were performed by the guidelines of the IACUC of the H. Lee Moffitt Cancer Center (that was approved by University of South Florida IACUC committee) using eight- to ten-week-old female nu/nu mice (Envigo). Mice had free access to water and food for the whole duration of the experiment. For MDA-MB-231 and MCF7 mono-tumor formation, 10 million cells were injected into cleared mammary fat pads as described previously (4). For mixed cultures, a 1:1 mixture of 5 million MDA-MB231 and 5 million MCF7 cells were injected. One week before cell injection, an estrogen pellet (0.72 mg slow-release, Innovative Research of America) was implanted to allow for the growth of ER-positive MCF7 tumors. Five weeks later, tumors were harvested.

## ACKNOWLEDGEMENTS

We thank Jeffrey West for assistance with software, we thank Chris Whelan and Dorothy Wallace for discussion and good ideas. Funding was provided by NIH/NCI 1U54CA193489-01A1, Cancer as a Complex Adaptive System, NIH/NCI U54 Supplement, The tumor-host evolutionary arms race, and R01 CA077575. This work was supported in part by the Analytic Microscopy Core Facility and the Tissue Core Facility at the H. Lee Moffitt Cancer Center & Research Institute; an NCI designated Comprehensive Cancer Center (P30-CA076292).

## AUTHOR CONTRIBUTIONS

All authors participated in the study ideas and the writing of the paper. M.D., A.I., A.R.F. and J.S.B designed the experiments. M.D., A.I. and A.R.F. collected the data. A.R.F., J.J.C. and J.S.B. performed the statistical analyses.

## References

1. Gillies RJ, Brown JS, Anderson AR, Gatenby RA. Eco-evolutionary causes and consequences of temporal changes in intratumoural blood flow. Nature Reviews Cancer. 2018;18(9):576–85.

2. Galon J, Costes A, Sanchez-Cabo F, Kirilovsky A, Mlecnik B, Lagorce-Pagès C, et al. Type, density, and location of immune cells within human colorectal tumors predict clinical outcome. Science. 2006;313(5795):1960–4

3. Galon J, Mlecnik B, Bindea G, Angell HK, Berger A, Lagorce C, et al. Towards the introduction of the ‘Immunoscore’in the classification of malignant tumours. The Journal of pathology. 2014;232(2):199–209.

4. Lloyd MC, Cunningham JJ, Bui MM, Gillies RJ, Brown JS, Gatenby RA. Darwinian dynamics of intratumoral heterogeneity: not solely random mutations but also variable environmental selection forces. Cancer research. 2016:canres.2962.015.

5. Ibrahim-Hashim A, Robertson-Tessi M, Enriquez-Navas PM, Damaghi M, Balagurunathan Y, Wojtkowiak JW, et al. Defining cancer subpopulations by adaptive strategies rather than molecular properties provides novel insights into intratumoral evolution. Cancer research. 2017;77(9):2242–54.

6. Gerlinger M, Rowan AJ, Horswell S, Larkin J, Endesfelder D, Gronroos E, et al. Intratumor heterogeneity and branched evolution revealed by multiregion sequencing. New England journal of medicine. 2012;366(10):883–92.

7. Polyak K. Heterogeneity in breast cancer. The Journal of clinical investigation. 2011;121(10):3786–8.

8. Perou CM, Sørlie T, Eisen MB, Van De Rijn M, Jeffrey SS, Rees CA, et al. Molecular portraits of human breast tumours. Nature. 2000;406(6797):747.

9. Robinson SP, Jordan VC. The paracrine stimulation of MCF-7 cells by MDA-MB-231 cells: possible role in antiestrogen failure. European Journal of Cancer and Clinical Oncology. 1989;25(3):493–7.

10. Lotka AJ. Contribution to the theory of periodic reactions. The Journal of Physical Chemistry. 1910;14(3):271–4.

11. Volterra V. Variazioni e fluttuazioni del numero d’individui in specie animali conviventi: C. Ferrari; 1927.

12. Gause G. The struggle for existence, 163 pp. Williams and Wilkins, Baltimore. 1934.

13. Gause G, Witt A. Behavior of Mixed Populations and the Problem of Natural Selection. The American Naturalist. 1935;69(725):569–609.

14. Barker J, Podger R. Interspecific competition between Drosophila melanogaster and Drosophila simulans: effects of larval density on viability, developmental period and adult body weight. Ecology. 1970;51(2):170–89.

15. Park T. Experimental studies of interspecies competition II. Temperature, humidity, and competition in two species of Tribolium. Physiological Zoology. 1954;27(3):177–238.

16. De Wit C, Van den Bergh J. Competition between herbage plants. Journal of Agricultural Science. 1965;13:212–21.

17. Kimmerling RJ, Prakadan SM, Gupta AJ, Calistri NL, Stevens MM, Olcum S, et al. Linking single-cell measurements of mass, growth rate, and gene expression. Genome biology. 2018;19(1):207.

18. Gerstein AC, Otto SP. Cryptic fitness advantage: diploids invade haploid populations despite lacking any apparent advantage as measured by standard fitness assays. PloS one. 2011;6(12):e26599.

19. Hart T, Chandrashekhar M, Aregger M, Steinhart Z, Brown KR, MacLeod G, et al. High-resolution CRISPR screens reveal fitness genes and genotype-specific cancer liabilities. Cell. 2015;163(6):1515–26.

20. Gallaher J, Brown J, Anderson ARA. The dynamic tumor ecosystem: how cell turnover and trade-offs affect cancer evolution. bioRxiv. 2018:270900.

21. Kaznatcheev A, Peacock J, Basanta D, Marusyk A, Scott JG. Fibroblasts and Alectinib switch the evolutionary games played by non-small cell lung cancer. bioRxiv. 2018:179259.

22. De Wit CT. On competition. Pudoc; 1960. Report No.: 0372-6223.

23. Rodríguez DJ. A method to study competition dynamics using de Wit replacement series experiments. Oikos. 1997;78(2):411–5.

24. Comşa Ş, Cimpean AM, Raica M. The story of MCF-7 breast cancer cell line: 40 years of experience in research. Anticancer research. 2015;35(6):3147–54.

25. Welsh J. Animal models for studying prevention and treatment of breast cancer. Animal models for the study of human disease: Elsevier; 2013. p. 997–1018.

26. Liu K, Newbury PA, Glicksberg BS, Zeng WZ, Paithankar S, Andrechek ER, et al. Evaluating cell lines as models for metastatic breast cancer through integrative analysis of genomic data. Nature communications. 2019;10(1):1–12.

27. Holliday DL, Speirs V. Choosing the right cell line for breast cancer research. Breast cancer research. 2011;13(4):215.

28. Neve RM, Chin K, Fridlyand J, Yeh J, Baehner FL, Fevr T, et al. A collection of breast cancer cell lines for the study of functionally distinct cancer subtypes. Cancer cell. 2006;10(6):515–27.

29. Kao J, Salari K, Bocanegra M, Choi Y-L, Girard L, Gandhi J, et al. Molecular profiling of breast cancer cell lines defines relevant tumor models and provides a resource for cancer gene discovery. PloS one. 2009;4(7):e6146.

30. Horwitz K, Costlow M, McGuire W. MCF-7: a human breast cancer cell line with estrogen, androgen, progesterone, and glucocorticoid receptors. Steroids. 1975;26(6):785–95.

31. Levenson AS, Jordan VC. MCF-7: the first hormone-responsive breast cancer cell line. Cancer research. 1997;57(15):3071–8.

32. Theodossiou TA, Ali M, Grigalavicius M, Grallert B, Dillard P, Schink KO, et al. Simultaneous defeat of MCF7 and MDA-MB-231 resistances by a hypericin PDT–tamoxifen hybrid therapy. NPJ breast cancer. 2019;5(1):1–10.

33. Sutherland RM, McCredie JA, Inch WR. Growth of multicell spheroids in tissue culture as a model of nodular carcinomas. Journal of the National Cancer Institute. 1971;46(1):113–20.

34. Benzekry S, Lamont C, Beheshti A, Tracz A, Ebos JM, Hlatky L, et al. Classical mathematical models for description and prediction of experimental tumor growth. PLoS computational biology. 2014;10(8):e1003800.

35. Marušić M, Bajzer Ž, Freyer J, Vuk-Pavlović S. Analysis of growth of multicellular tumour spheroids by mathematical models. Cell proliferation. 1994;27(2):73–94.

36. Kucharavy D, De Guio R. Application of logistic growth curve. Procedia engineering. 2015;131:280–90.

37. Vaghi C, Rodallec A, Fanciullino R, Ciccolini J, Mochel JP, Mastri M, Poignard C, Ebos JM, Benzekry S. Population modeling of tumor growth curves and the reduced Gompertz model improve prediction of the age of experimental tumors. PLoS computational biology. 2020;16(2):e1007178.

38. Monod J. The growth of bacterial cultures. Annual review of microbiology. 1949;3(1):371–94.

39. Liancourt P, Callaway RM, Michalet R. Stress tolerance and competitive-response ability determine the outcome of biotic interactions. Ecology. 2005;86(6):1611–8.

40. Douda J, Doudová J, Hulík J, Havrdová A, Boublík K. Reduced competition enhances community temporal stability under conditions of increasing environmental stress. Ecology. 2018;99(10):2207–16.

41. Igaki, T., Pastor-Pareja, J.C., Aonuma, H., Miura, M. and Xu, T., 2009. Intrinsic tumor suppression and epithelial maintenance by endocytic activation of Eiger/TNF signaling in Drosophila. Developmental cell, 16(3), pp.458–465.

42. Ballesteros-Arias, L., Saavedra, V. and Morata, G., 2014. Cell competition may function either as tumour-suppressing or as tumour-stimulating factor in Drosophila. Oncogene, 33(35), pp.4377–4384.

43. Nishikawa, S., Takamatsu, A., Ohsawa, S. and Igaki, T., 2016. Mathematical model for cell competition: Predator–prey interactions at the interface between two groups of cells in monolayer tissue. Journal of theoretical biology, 404, pp.40–50.

44. Michl, Johanna, Kyung Chan Park, and Pawel Swietach. “Evidence-based guidelines for controlling pH in mammalian live-cell culture systems.” Communications biology 2, no. 1 (2019): 1–12.

45. Amend SR, Gatenby RA, Pienta KJ, Brown JS. Cancer foraging ecology: Diet choice, patch use, and habitat selection of cancer cells. Current Pathobiology Reports. 2018;6(4):209–18.

46. Brown JS. Coexistence on a seasonal resource. The American Naturalist. 1989;133(2):168–82.

47. Chesson P, Gebauer RL, Schwinning S, Huntly N, Wiegand K, Ernest MS, et al. Resource pulses, species interactions, and diversity maintenance in arid and semi-arid environments. Oecologia. 2004;141(2):236–53.

48. Wells A, Grahovac J, Wheeler S, Ma B, Lauffenburger D. Targeting tumor cell motility as a strategy against invasion and metastasis. Trends in pharmacological sciences. 2013;34(5):283–9.

49. Hagy HM, Kaminski RM. Determination of foraging thresholds and effects of application on energetic carrying capacity for waterfowl. PLoS One. 2015;10(3).

50. Richards SA, Nisbet RM, Wilson WG, Possingham HP. Grazers and diggers: exploitation competition and coexistence among foragers with different feeding strategies on a single resource. The American Naturalist. 2000;155(2):266–79.

51. Taylor TB, Wass AV, Johnson LJ, Dash P. Resource competition promotes tumour expansion in experimentally evolved cancer. BMC evolutionary biology. 2017;17(1):268.

52. Beaupain R. A method for three-dimensional coculture of cancer cells combined to any other type of cells maintained organotypically. Methods in cell science. 1999;21(1):25–30.

53. Hirschhaeuser F, Menne H, Dittfeld C, West J, Mueller-Klieser W, Kunz-Schughart LA. Multicellular tumor spheroids: an underestimated tool is catching up again. Journal of biotechnology. 2010;148(1):3–15.

54. Weiswald L-B, Bellet D, Dangles-Marie V. Spherical cancer models in tumor biology. Neoplasia. 2015;17(1):1–15.

55. Sutherland RM. Importance of critical metabolites and cellular interactions in the biology of microregions of tumors. Cancer. 1986;58(8):1668–80.

56. Sutherland RM. Cell and environment interactions in tumor microregions: the multicell spheroid model. Science. 1988;240(4849):177–84.

57. Mueller-Klieser W, Freyer J, Sutherland R. Influence of glucose and oxygen supply conditions on the oxygenation of multicellular spheroids. British journal of cancer. 1986;53(3):345.

58. Tofilon PJ, Arundel CM, Deen DF. Response to BCNU of spheroids grown from mixtures of drug-sensitive and drug-resistant cells. Cancer chemotherapy and pharmacology. 1987;20(2):89–95.

59. Gatenby RA, Silva AS, Gillies RJ, Frieden BR. Adaptive therapy. Cancer research. 2009;69(11):4894–903.

60. Gallaher JA, Enriquez-Navas PM, Luddy KA, Gatenby RA, Anderson AR. Spatial heterogeneity and evolutionary dynamics modulate time to recurrence in continuous and adaptive cancer therapies. Cancer research. 2018;78(8):2127–39.

61. Zhang J, Cunningham JJ, Brown JS, Gatenby RA. Integrating evolutionary dynamics into treatment of metastatic castrate-resistant prostate cancer. Nature communications. 2017;8(1):1816.

62. Rasband, W.S., ImageJ, U. S. National Institutes of Health, Bethesda, Maryland, USA, https://imagej.nih.gov/ij/, 1997-2018.

