## Supplementary Material for "Frequency-dependent interactions determine outcome of competition between two breast cancer cell lines"

| Variable | Type III Sums of Squares | Degrees Freedom | Mean Squares | F-Ratio | P-Value |
|---|---|---|---|---|---|
| **MCF-7 Instantaneous Growth Rates** | | | | | |
| **Glucose Concentration (GC)** | 18.528 | 1 | 18.528 | 62.389 | < 0.00 |
| **pH** | 0.014 | 1 | 0.014 | 0.047 | 0.829 |
| **Starting Frequency (f)** | 21.514 | 4 | 5.379 | 18.111 | < 0.000 |
| **pH x GC** | 5.270 | 1 | 5.270 | 17.747 | < 0.000 |
| **f x GC** | 0.812 | 4 | 0.203 | 0.684 | 0.606 |
| **pH x f** | 1.755 | 4 | 0.439 | 1.477 | 0.220 |
| **Error** | 18.710 | 63 | 0.297 | | |
| **MDA-MB-231 Instantaneous Growth Rates** | | | | | |
| **Glucose Concentration (GC)** | 0.038 | 1 | 0.038 | 13.957 | < 0.00 |
| **pH** | 0.172 | 1 | 0.172 | 62.881 | < 0.00 |
| **Starting Frequency (f)** | 1.157 | 4 | 0.289 | 105.748 | < 0.00 |
| **pH x GC** | 0.002 | 1 | 0.002 | 0.761 | 0.386 |
| **f x GC** | 0.002 | 4 | 0.001 | 0.224 | 0.924 |
| **pH x f** | 0.011 | 4 | 0.003 | 0.971 | 0.430 |
| **Error** | 0.175 | 64 | 0.003 | | |
| **Difference Between MCF-7 and MDA-MB-231 Growth Rates** | | | | | |
| **Glucose Concentration (GC)** | 19.473 | 1 | 19.473 | 57.178 | < 0.00 |
| **pH** | 0.013 | 1 | 0.013 | 0.038 | 0.846 |
| **Starting Frequency (f)** | 1.680 | 3 | 0.560 | 1.644 | 0.191 |
| **pH x GC** | 4.896 | 1 | 4.896 | 14.377 | < 0.000 |
| **f x GC** | 0.281 | 3 | 0.094 | 0.275 | 0.843 |
| **pH x f** | 0.999 | 3 | 0.333 | 0.977 | 0.411 |
| **Error** | 17.369 | 51 | 0.341 | | |

**Supplementary table 1:** ANOVA results to determine statistical significance of culture conditions on growth rate for experiment with high seeding density (20,000 cells) and glucose concentrations of 0 g/L and 2 g/L.

| MCF-7 Instantaneous Growth Rates | | | | | |
|---|---|---|---|---|---|
| **Variable** | **Type III Sums of Squares** | **Degrees Freedom** | **Mean Squares** | **F-Ratio** | **P-Value** |
| **Glucose Concentration (GC)** | 0.018 | 1 | 0.018 | 0.138 | 0.711 |
| **pH** | 1.126 | 1 | 1.126 | 8.852 | 0.004 |
| **Starting Frequency (f)** | 26.713 | 4 | 6.678 | 52.525 | < 0.000 |
| **pH x GC** | 0.953 | 1 | 0.953 | 7.499 | 0.008 |
| **f x GC** | 0.298 | 4 | 0.075 | 0.586 | 0.674 |
| **pH x f** | 1.307 | 4 | 0.327 | 2.570 | 0.046 |
| **Error** | 8.137 | 64 | 0.127 | | |
| **MDA-MB-231 Instantaneous Growth Rates** | | | | | |
| **Glucose Concentration (GC)** | 0.009 | 1 | 0.009 | 4.547 | 0.037 |
| **pH** | 0.111 | 1 | 0.111 | 53.146 | < 0.000 |
| **Starting Frequency (f)** | 0.964 | 4 | 0.241 | 115.872 | < 0.000 |
| **pH x GC** | 0.006 | 1 | 0.006 | 2.727 | 0.104 |
| **f x GC** | 0.009 | 4 | 0.002 | 1.115 | 0.357 |
| **pH x f** | 0.041 | 4 | 0.010 | 4.987 | 0.001 |
| **Error** | 0.133 | 64 | 0.002 | | |
| **Difference Between MCF-7 and MDA-MB-231 Growth Rates** | | | | | |
| **Glucose Concentration (GC)** | 0.004 | 1 | 0.004 | 0.024 | 0.877 |
| **pH** | 1.392 | 1 | 1.392 | 9.390 | 0.003 |
| **Starting Frequency (f)** | 10.427 | 3 | 3.476 | 23.445 | < 0.000 |
| **pH x GC** | 1.049 | 1 | 1.049 | 7.075 | 0.010 |
| **f x GC** | 0.186 | 3 | 1.049 | 0.419 | 0.740 |
| **pH x f** | 1.236 | 3 | 0.412 | 2.778 | 0.050 |
| **Error** | 7.561 | 51 | 0.148 | | |

**Supplementary table 2:** ANOVA results to determine statistical significance of culture conditions on growth rate for experiment with high seeding density (20,000 cells) and glucose concentrations of 1 g/L and 4.5 g/L.

| MCF-7 Instantaneous Growth Rates +glutamine | | | | | |
|---|---|---|---|---|---|
| **Variable** | **Type III Sums of Squares** | **Degrees Freedom** | **Mean Squares** | **F-Ratio** | **P-Value** |
| **Glucose Concentration (GC)** | 4.955 | 4 | 1.239 | 3.640 | 0.011 |
| **pH** | 0.538 | 1 | 0.538 | 1.580 | 0.214 |
| **Starting Frequency (f)** | 0.216 | 1 | 0.216 | 0.636 | 0.429 |
| **pH x GC** | 1.504 | 4 | 0.376 | 1.104 | 0.364 |
| **f x GC** | 0.784 | 4 | 0.196 | 0.576 | 0.681 |
| **pH x f** | 0.443 | 1 | 0.443 | 1.301 | 0.259 |
| **Error** | 18.721 | 55 | 0.340 | | |
| **MDA-MB-231 Instantaneous Growth Rates +glutamine** | | | | | |
| **Glucose Concentration (GC)** | 0.074 | 1 | 0.074 | 4.283 | 0.043 |
| **pH** | 0.127 | 1 | 0.127 | 7.318 | 0.009 |
| **Starting Frequency (f)** | 0.904 | 4 | 0.226 | 13.004 | < 0.000 |
| **pH x GC** | 0.006 | 1 | 0.006 | 0.367 | 0.547 |
| **f x GC** | 0.014 | 4 | 0.004 | 0.204 | 0.935 |
| **pH x f** | 0.116 | 4 | 0.029 | 1.672 | 0.167 |
| **Error** | 1.113 | 64 | 0.017 | | |
| **MCF-7 Instantaneous Growth Rates -glutamine** | | | | | |
| **Variable** | **Type III Sums of Squares** | **Degrees Freedom** | **Mean Squares** | **F-Ratio** | **P-Value** |
| **Glucose Concentration (GC)** | 1.655 | 1 | 1.655 | 3.177 | 0.080 |
| **pH** | 5.023 | 1 | 5.023 | 9.641 | 0.003 |
| **Starting Frequency (f)** | 12.791 | 4 | 3.198 | 6.138 | < 0.000 |
| **pH x GC** | 2.309 | 1 | 2.309 | 4.432 | 0.039 |
| **f x GC** | 4.895 | 4 | 1.224 | 2.349 | 0.064 |
| **pH x f** | 7.648 | 4 | 1.912 | 3.670 | 0.010 |
| **Error** | 31.262 | 60 | 0.521 | | |
| **MDA-MB-231 Instantaneous Growth Rates -glutamine** | | | | | |
| **Glucose Concentration (GC)** | 0.016 | 1 | 0.016 | 1.116 | 0.295 |
| **pH** | 0.113 | 1 | 0.113 | 7.968 | 0.006 |
| **Starting Frequency (f)** | 0.930 | 4 | 0.233 | 16.358 | < 0.000 |
| **pH x GC** | 0.026 | 1 | 0.026 | 1.794 | 0.185 |
| **f x GC** | 0.098 | 4 | 0.025 | 1.731 | 0.154 |
| **pH x f** | 0.146 | 4 | 0.036 | 2.560 | 0.047 |
| **Error** | 0.910 | 64 | 0.014 | | |

**Supplementary table 3:** ANOVA results to determine statistical significance of culture conditions on growth rate for experiment with low seeding density (10,000 cells) and glucose concentrations of 0 g/L and 4.5 g/L.

| MCF-7 Carrying Capacity | | | | | |
|---|---|---|---|---|---|
| **Variable** | **Type III Sums of Squares** | **Degrees Freedom** | **Mean Squares** | **F-Ratio** | **P-Value** |
| **Glucose Concentration (GC)** | 1.503 | 1 | 1.503 | 0.008 | 0.930 |
| **pH** | 195.016 | 1 | 195.016 | 1.047 | 0.328 |
| **pH x GC** | 1,384.361 | 1 | 1,384.361 | 7.435 | 0.020 |
| **Error** | 2,048.017 | 11 | 186.183 | | |
| **MDA-MB-231 Carrying Capacity** | | | | | |
| **Glucose Concentration (GC)** | 1,116.013 | 1 | 1,116.013 | 610.634 | < 0.00 |
| **pH** | 1,237.514 | 1 | 1,237.514 | 677.114 | < 0.00 |
| **pH x GC** | 972.488 | 1 | 972.488 | 532.103 | < 0.000 |
| **Error** | 116.968 | 64 | 1.828 | | |

| MCF-7 Carrying Capacity | | | | | |
|---|---|---|---|---|---|
| **Variable** | **Type III Sums of Squares** | **Degrees Freedom** | **Mean Squares** | **F-Ratio** | **P-Value** |
| **Glucose Concentration (GC)** | 18.431 | 1 | 18.431 | 0.400 | 0.539 |
| **pH** | 593.962 | 1 | 593.962 | 12.875 | 0.004 |
| **pH x GC** | 398.561 | 1 | 398.561 | 8.639 | 0.012 |
| **Error** | 553.593 | 12 | 46.133 | | |
| **MDA-MB-231 Carrying Capacity** | | | | | |
| **Glucose Concentration (GC)** | 181.817 | 1 | 181.817 | 67.181 | < 0.000 |
| **pH** | 66.078 | 1 | 66.078 | 24.416 | < 0.000 |
| **pH x GC** | 71.761 | 1 | 71.761 | 26.516 | < 0.000 |
| **Error** | 205.683 | 76 | 2.706 | | |

**Supplementary table 4:** ANOVA results to determine statistical significance of culture conditions on carrying capacity for experiment with high seeding density (20,000 cells) and glucose concentrations of 0 g/L and 2 g/L in top table and 1 g/L and 4.5 g/L in lower table.

| MCF-7 Carrying Capacity +glutamine | | | | | |
|---|---|---|---|---|---|
| Variable | Type III Sums of Squares | Degrees Freedom | Mean Squares | F-Ratio | P-Value |
| Glucose Concentration (GC) | 371.403 | 1 | 371.403 | 8.314 | 0.005 |
| pH | 270.455 | 1 | 270.455 | 6.054 | 0.017 |
| Starting Frequency (f) | 8,201.115 | 4 | 2,050.279 | 45.896 | < 0.000 |
| pH x GC | 0.109 | 1 | 0.109 | 0.002 | 0.961 |
| f x GC | 481.061 | 4 | 120.265 | 2.692 | 0.039 |
| pH x f | 172.672 | 4 | 43.168 | 0.966 | 0.432 |
| Error | 2,858.997 | 64 | 44.672 | | |
| **MDA-MB-231 Carrying Capacity + glutamine** | | | | | |
| Glucose Concentration (GC) | 0.856 | 1 | 0.856 | 1.784 | 0.186 |
| pH | 125.666 | 1 | 125.666 | 261.952 | < 0.000 |
| Starting Frequency (f) | 220.961 | 4 | 55.240 | 115.149 | < 0.000 |
| pH x GC | 1.260 | 1 | 1.260 | 2.626 | 0.110 |
| f x GC | 7.240 | 4 | 1.810 | 3.773 | 0.008 |
| pH x f | 57.634 | 4 | 14.408 | 30.034 | < 0.000 |
| Error | 30.703 | 64 | 0.480 | | |

| MCF-7 Carrying Capacity -glutamine | | | | | |
|---|---|---|---|---|---|
| Variable | Type III Sums of Squares | Degrees Freedom | Mean Squares | F-Ratio | P-Value |
| Glucose Concentration (GC) | 1,419,397.848 | 1 | 1,419,397.848 | 3.890 | 0.053 |
| pH | 1,941,483.025 | 1 | 1,941,483.025 | 5.321 | 0.024 |
| Starting Frequency (f) | 1.726E+008 | 4 | 43,154,454.625 | 118.278 | < 0.000 |
| pH x GC | 1,291,221.858 | 1 | 1,291,221.858 | 3.539 | 0.064 |
| f x GC | 686,257.624 | 4 | 171,564.406 | 0.470 | 0.757 |
| pH x f | 16,814,748.188 | 4 | 4,203,687.047 | 11.521 | < 0.000 |
| Error | 23,350,855.237 | 64 | 364,857.113 | | |
| **MDA-MB-231 Carrying Capacity -glutamine** | | | | | |
| Glucose Concentration (GC) | 47,425.034 | 1 | 47,425.034 | 7.808 | 0.007 |
| pH | 22,960.408 | 1 | 22,960.408 | 3.780 | 0.056 |
| Starting Frequency (f) | 1,143,424.578 | 4 | 285,856.144 | 47.060 | < 0.000 |
| pH x GC | 78,851.808 | 1 | 78,851.808 | 12.981 | 0.001 |
| f x GC | 156,395.360 | 4 | 39,098.840 | 6.437 | < 0.000 |
| pH x f | 26,175.583 | 4 | 6,543.896 | 1.077 | 0.375 |
| Error | 388,752.068 | 64 | 6,074.251 | | |

**Supplementary table 5:** ANOVA results to determine statistical significance of culture conditions on carrying capacity for experiment with low seeding density (10,000 cells) and glucose concentrations of 0 g/L and 4.5 g/L.

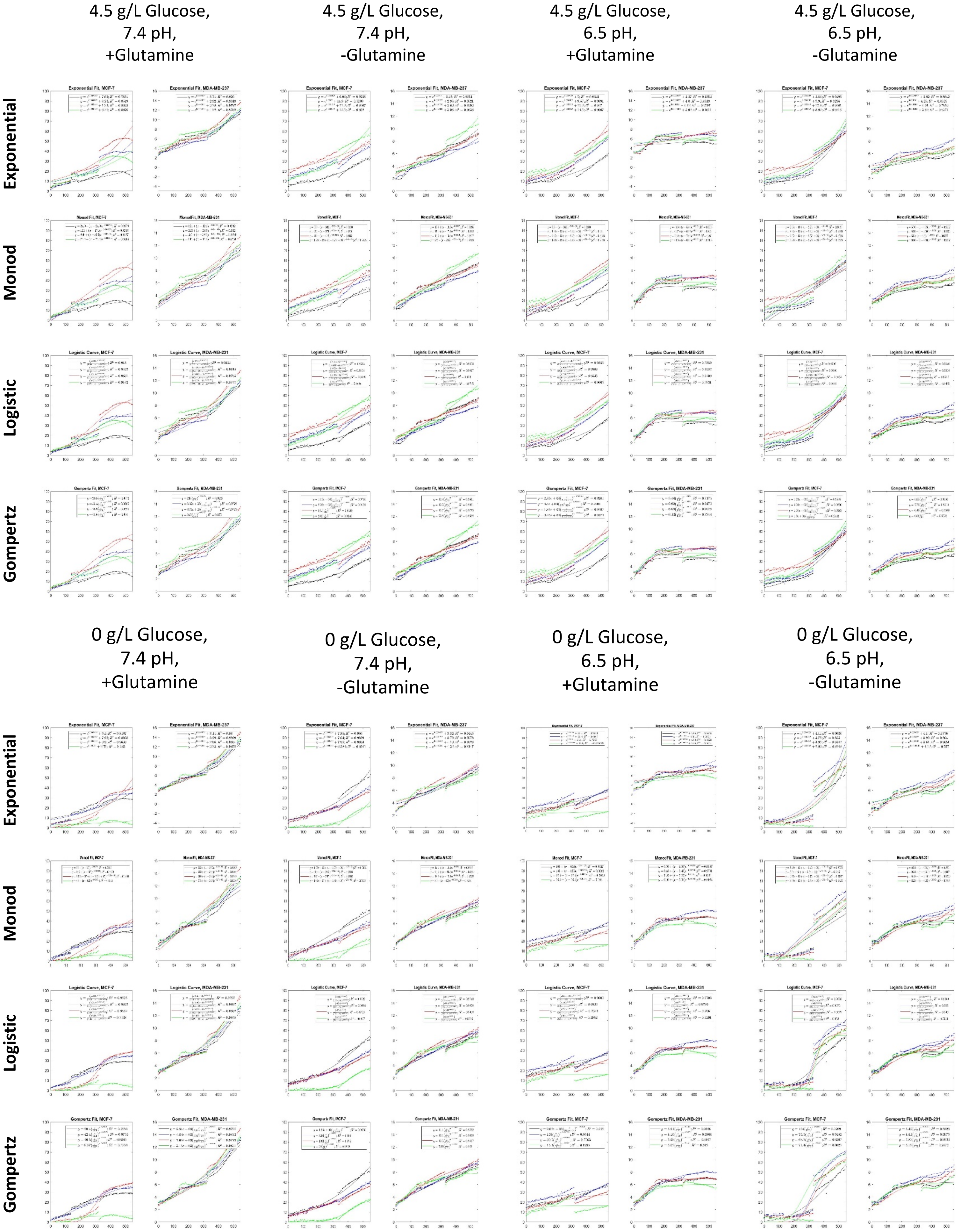


**Supplementary Figure 1:** Fits of the exponential, Monod-like, logistic and Gompertz growth models to monoculture spheroid data. Fitting was performed using nonlinear constrained optimization.

| Variable | Type III Sums of Squares | Degrees Freedom | Mean Squares | F-Ratio | P-Value |
|---|---|---|---|---|---|
| **Adjusted $R^2$** | | | | | |
| **Model** | 0.430 | 3 | 0.143 | 9.966 | < 0.000 |
| **Culture Condition** | 1.013 | 7 | 0.145 | 10.074 | < 0.000 |
| **Cell Type** | 0.025 | 1 | 0.025 | 1.726 | 0.190 |
| **Culture Condition x Model** | 0.272 | 21 | 0.013 | 0.901 | 0.589 |
| **Cell Type x Model** | 0.154 | 3 | 0.051 | 3.568 | 0.015 |
| **Cell Type x Culture Condition** | 0.990 | 7 | 0.141 | 9.845 | < 0.000 |
| **Error** | 3.060 | 213 | 0.014 | | |
| **RMSE** | | | | | |
| **Model** | 280.951 | 3 | 93.650 | 5.894 | 0.001 |
| **Culture Condition** | 2,670.428 | 7 | 381.490 | 24.009 | < 0.000 |
| **Cell Type** | 17,782.592 | 1 | 17,782.592 | 1,119.135 | < 0.000 |
| **Culture Condition x Model** | 57.078 | 21 | 2.718 | 0.171 | 1.000 |
| **Cell Type x Model** | 127.607 | 3 | 42.536 | 2.677 | 0.048 |
| **Cell Type x Culture Condition** | 3,175.509 | 7 | 453.644 | 28.550 | < 0.000 |
| **Error** | 3,384.481 | 213 | 15.890 | | |

**Supplementary table 6:** Results of ANOVA comparing quality of fit measurements (Adjusted $R^2$ and RMSE) for all four growth models.

| MCF-7 Slope of Carrying Capacity +glutamine | | | | | |
|---|---|---|---|---|---|
| **Variable** | **Type III Sums of Squares** | **Degrees Freedom** | **Mean Squares** | **F-Ratio** | **P-Value** |
| **Glucose Concentration (GC)** | 0.000 | 1 | 0.000 | 0.099 | 0.754 |
| **pH** | 0.030 | 1 | 0.030 | 86.905 | < 0.000 |
| **Starting Frequency (f)** | 0.038 | 4 | 0.009 | 27.563 | < 0.000 |
| **pH x GC** | 0.003 | 1 | 0.003 | 7.412 | 0.008 |
| **f x GC** | 0.000 | 4 | 0.000 | 0.298 | 0.878 |
| **pH x f** | 0.026 | 4 | 0.006 | 18.994 | < 0.000 |
| **Error** | 0.022 | 64 | 0.000 | | |
| **MDA-MB-231 Slope of Carrying Capacity +glutamine** | | | | | |
| **Glucose Concentration (GC)** | 0.000 | 1 | 0.000 | 16.406 | < 0.000 |
| **pH** | 0.003 | 1 | 0.003 | 358.724 | < 0.000 |
| **Starting Frequency (f)** | 0.001 | 4 | 0.000 | 19.839 | < 0.000 |
| **pH x GC** | 0.000 | 1 | 0.000 | 1.528 | 0.221 |
| **f x GC** | 0.000 | 4 | 0.000 | 1.084 | 0.372 |
| **pH x f** | 0.002 | 4 | 0.000 | 56.042 | < 0.000 |
| **Error** | 0.000 | 64 | 0.000 | | |
| **MCF-7 Slope of Carrying Capacity -glutamine** | | | | | |
| **Variable** | **Type III Sums of Squares** | **Degrees Freedom** | **Mean Squares** | **F-Ratio** | **P-Value** |
| **Glucose Concentration (GC)** | 1.019 | 1 | 1.019 | 0.837 | 0.364 |
| **pH** | 3.316 | 1 | 3.316 | 2.723 | 0.104 |
| **Starting Frequency (f)** | 1,287.534 | 4 | 321.884 | 264.291 | < 0.000 |
| **pH x GC** | 0.223 | 1 | 0.223 | 0.183 | 0.670 |
| **f x GC** | 10.263 | 4 | 2.566 | 2.107 | 0.090 |
| **pH x f** | 100.048 | 4 | 25.012 | 20.537 | < 0.000 |
| **Error** | 77.946 | 64 | 1.218 | | |
| **MDA-MB-231 Slope of Carrying Capacity −glutamine** | | | | | |
| **Glucose Concentration (GC)** | 0.618 | 1 | 0.618 | 9.652 | 0.003 |
| **pH** | 0.139 | 1 | 0.139 | 2.179 | 0.145 |
| **Starting Frequency (f)** | 1.793 | 4 | 0.448 | 7.005 | < 0.000 |
| **pH x GC** | 1.151 | 1 | 1.151 | 17.982 | < 0.000 |
| **f x GC** | 0.234 | 4 | 0.058 | 0.912 | 0.462 |
| **pH x f** | 1.646 | 4 | 0.412 | 6.430 | < 0.000 |
| **Error** | 4.096 | 64 | 0.064 | | |

**Supplementary table 7:** ANOVA comparing the final slopes of the low seeding density spheroids. Carrying capacity values were determined using the monoculture spheroids for MCF-7 and all spheroids for MDA-MB-231.
